# Instability with a purpose: how the visual brain makes decisions in a volatile world

**DOI:** 10.1101/2020.06.09.142497

**Authors:** Robin Cao, Alexander Pastukhov, Stepan Aleshin, Maurizio Mattia, Jochen Braun

## Abstract

In ambiguous or conflicting sensory situations, perception is often ‘multistable’ in that it changes abruptly at irregular intervals, shifting perpetually between distinct alternatives. Intriguingly, the interval statistics of these alternations exhibits quasi-universal characteristics, suggesting a general mechanism. Here we show that the stereotypical features of multistable perception, exemplified by binocular rivalry, are reproduced in detail by a hierarchical dynamics operating out of equilibrium. Its constitutive elements are discretely stochastic and idealize the metastability of cortical networks. Independent elements accumulate visual evidence at one level, while groups of coupled elements compete for dominance at another level. As soon as one group dominates perception, feedback inhibition suppresses supporting evidence. This mechanism is corroborated compellingly by unexpected serial dependencies of perceptual alternations. Moreover, it satisfies normative constraints of continuous decision-making. We conclude that multistable perception reflects decision-making in a volatile world: integrating evidence over space and time, choosing categorically between hypotheses, while concurrently evaluating alternatives.

## INTRODUCTION

In deducing the likely physical causes of sensations, perception goes beyond the immediate sensory evidence and draws heavily on context and prior experience [5, 44, 100, 118]. Numerous illusions in visual, auditory, and tactile perception – all subjectively compelling, but objectively false – attest to this extrapolation beyond the evidence. In natural settings, perception explores alternative plausible causes of sensory evidence by active repositioning of sensors [“active perception”, 78, 85, 129]. In general, perception is thought to actively select plausible explanatory hypotheses, to predict the sensory evidence expected for each hypothesis from prior experience, and to compare the observed sensory evidence at multiple levels of scale or abstraction [“analysis by synthesis”, “predictive coding”, “hierarchical Bayesian inference”, 86, 91, 97, 130]. Active inference engages the entire hierarchy of sensory cortical areas, including both feedforward and feedback projections [4, 40, 60, 87, 107].

The dynamics of active inference becomes experimentally observable when perceptual illusions are ‘multistable’ [65]. In numerous ambiguous or conflicting situations, phenomenal experience switches at irregular intervals between discrete alternatives, even though the sensory scene is stable [3, 83, 95, 96, 102, 105, 126]. Multistable illusions are enormously diverse, involving visibility or audibility, perceptual grouping, visual depth or motion, and many kinds of sensory scenes, from schematic to naturalistic. Average switching rates differ greatly and range over at least two orders of magnitude [19], depending on sensory scene, perceptual grouping [53, 112, 125], continuous or intermittent presentation [64, 72], attentional condition [88], individual observer [16, 32, 89], and many other factors.

In spite of this diversity, the stochastic properties of multistable phenomena appear to be quasi-universal, suggesting that the underlying mechanisms may be general. Firstly, average dominance duration depends in a characteristic and counter-intuitive manner on the strength of dominant and suppressed evidence [“Levelt’s propositions I–IV”, 14, 15, 48, 51, 66, 80]. Secondly, the statistical distribution of dominance durations shows a stereotypical shape, namely a Gamma distribution with shape parameter *r* ≃ 3−4 [“scaling property”, 8, 12, 13, 19, 28, 32, 39, 82, 88, 119]. Thirdly, the durations of successive dominance periods are correlated positively, over at least two or three periods [32, 39, 114, 119].

Here we show that these quasi-universal characteristics are comprehensively and quantitatively reproduced, indeed guaranteed, by an interacting hierarchy of birth-death processes. operating out of equilibrium. While the proposed mechanism combines key features of previous models, it far surpasses their explanatory power.

Several possible mechanisms have been proposed for perceptual dominance, the triggering of reversals, and the stochastic timing of reversals. That a single, coherent interpretation typically dominates phenomenal experience is thought to reflect competition (explicit or implicit) at the level of explanatory hypotheses [*e.g*., 27], sensory inputs [*e.g*., 63], or both [*e.g*., 127]. That a dominant interpretation is periodically supplanted by a distinct alternative has been attributed to fatigue processes [*e.g*., neural adaptation, synaptic depression, 59], spontaneous fluctuations [‘noise’, *e.g*., 50, 128], stochastic sampling [*e.g*., 104], or combinations of these [*e.g*., adaptation and noise, 89, 106, 108]. The characteristic stochasticity (Gamma-like distribution) of dominance durations has been attributed to Poisson counting processes [*e.g*., birth-death processes, 19, 41, 111] or stochastic accumulation of discrete samples [82, 104, 110, 124].

‘Dynamical’ models combining competition, adaptation, and noise capture well the characteristic dependence of dominance durations on input strength (“Levelt’s propositions”) [2, 59, 128], especially when inputs are normalized [23, 79, 80], and when the dynamics emphasize noise [89, 106, 108]. However, such models do not preserve a Gamma-like distribution over the full range of input strengths [19, 23]. On the other hand, ‘sampling’ models based on discrete random processes produce Gamma-like distributions, [19, 82, 104, 110, 111, 124], but fail to reproduce the dependence on input strength. Neither type of model accounts for the sequential dependence of dominance durations [59].

Here we reconcile ‘dynamical’ and ‘sampling’ approaches to multistable perception, extending an earlier effort [41]. Importantly, every part of the proposed mechanism is justified normatively in that it serves to optimize perceptual choices in a general behavioural situation, namely, continuous inference in uncertain and volatile environments [9, 117]. We propose that sensory inputs are represented by birth-death processes, in order to accumulate sensory information over time and in a format suited for Bayesian inference [70, 93]. Further, we suggest that explanatory hypotheses are evaluated competitively, with a hypothesis attaining dominance (over phenomenal experience) when its support exceeds the alternatives by a certain finite amount, consistent with optimal decision making between multiple alternatives [9]. Finally, we assume that a dominant hypothesis suppresses its supporting evidence, as required by ‘predictive coding’ implementations of hierarchical Bayesian inference [46, 90, 97]. In contrast to many previous models, we do not require a mechanism of fatigue, adaptation, or decay.

Based on these assumptions, the proposed mechanism reproduces dependence on input strength, as well distribution of dominance durations and positive sequential dependence. Additionally, it predicts novel and unsuspected dynamical features confirmed by experiment.

## RESULTS

Next we introduce each component of the mechanism with its normative justification and describe the reversal dynamics resulting from the interaction of all components. Further below, we compare model predictions with multistable perception of human observers, specifically, the dominance statistics of binocular rivalry at various combinations of left- and right-eye contrasts (**Fig. 1a**).

**Figure 1.**
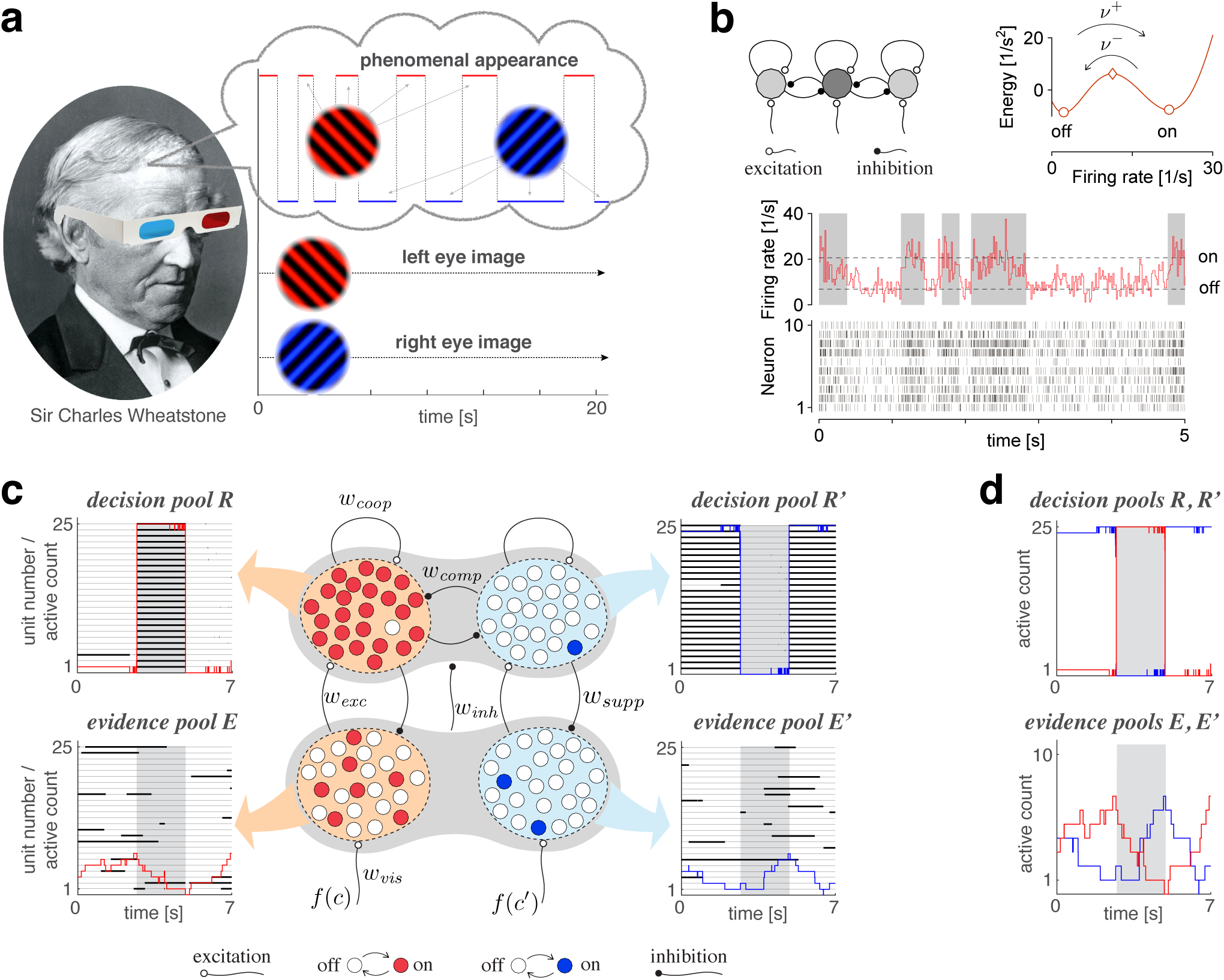
Proposed mechanism of binocular rivalry. **(a)** When the left and right eyes see incompatible images, phenomenal appearance reverses at irregular intervals, sometimes being dominated by one image and sometimes by the other (grey and white regions). Sir Charles Wheatstone studied this multistable percept with a mirror stereoscope (not as shown!). **(b)** Spiking neural network implementation of a ‘local attractor’. An assembly of 150 neurons (schematic, dark gray circle) interacts competitively with multiple other assemblies (light gray circles). Population activity of the assembly explores an effective energy landscape (right) with two distinct steady-states (circles), separated by a ridge (diamond). Driven by noise, activity transitions occasionally between ‘on’ and ‘off’ states (bottom), with transition rates *ν*^±^ depending sensitively on external input to the assembly (not shown). Here, *ν*^+^=*ν*^−^≈1 *Hz*. Spike raster shows 10 representative neurons. **(c)** Nested attractor dynamics (schematic) which quantitatively reproduces the dynamics of binocular rivalry. Independently bistable variables (‘local attractors’, small circles) respond probabilistically to input, transitioning stochastically between on- and off-states (red and white, respectively). The entire system comprises four pools, with 25 variables each, linked by excitatory and inhibitory projections. Phenomenal appearance is decided by competition between decision pools *R* and *R*′ forming ‘non-local attractors’ (cross-inhibition *w*_*comp*_ and self-excitation *w*_*coop*_). Visual input *c* and *c*′ accumulates, respectively, in evidence pools *E* and *E*′ and propagates to decision pools (feedforward selective excitation *w*_*exc*_ and indiscriminate inhibition *w*_*inh*_). Decision pools suppress associated evidence pools (feedback selective suppression *w*_*supp*_). **(d)** Representative example of model dynamics. The time-course of the number of active variables (active count) is shown for decision pools (top) and evidence pools (bottom), representing the left eye (red traces) and the right eye image (blue traces). Left and right pools are shown both separately (left and middle columns, respectively) and together (right column). The state of individual variables (black horizontal traces in left and middle columns) and of perceptual dominance (grey and white regions) are also shown. In decision pools, almost all variables become active (thick trace) or inactive (thin trace) simultaneously. In evidence pools, only a small fraction of variables is active at any given time.

### Hierarchical dynamics

#### Bistable assemblies: ‘local attractors’

As operative units of sensory representation, we postulate neuronal assemblies with bistable ‘attractor’ dynamics. Effectively, assembly activity moves in an energy landscape with two distinct quasi-stable states – dubbed ‘on’ and ‘off’ – separated by a ridge (**Fig. 1b**). Driven by noise, assembly activity mostly remains near one quasi-stable state (‘on’ or ‘off’), but occasionally ‘escapes’ to the other state [29, 45, 47, 55, 67].

An important feature of ‘attractor’ dynamics is that the energy of quasi-stable states depends sensitively on external input. Net positive input destabilizes (*i.e*., raises the potential of) the ‘off’ state and stabilizes (*i.e*., lowers the potential of) the ‘on’ state. Transition rates *ν*^±^ depend approximately *exponentially* on the height of the energy ridge (‘activation energy’).

**Figure 1b** illustrates ‘attractor’ dynamics for an assembly of 150 spiking neurons with activity levels of approximately 7 *Hz* and 21 *Hz* per neuron in the ‘off’ and ‘on’ states, respectively. Further details are given in the supplementary text (**Metastable population dynamics** and **Fig. 8**). Note that our model is independent of network details and relies only on an idealized description.

**Figure 2.**
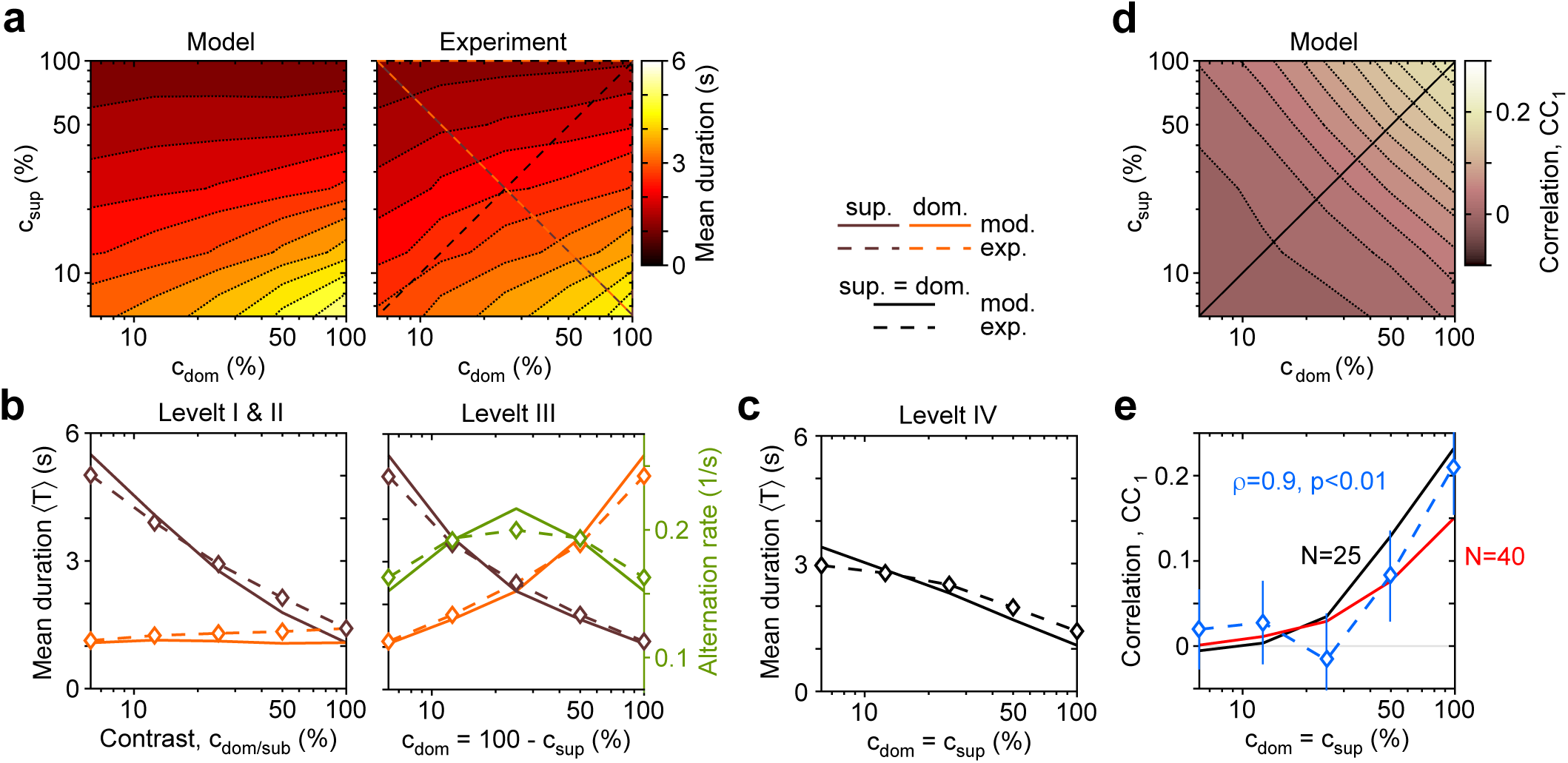
Dependence of mean dominance duration on dominant and suppressed image contrast (“Levelt’s propositions”). **(a)** Mean dominance duration ⟨*T*⟩ (color scale), as a function of dominant contrast *c*_*dom*_ and suppressed contrast *c*_*sup*_, in model (left) and experiment (right). **(b)** Model prediction (solid traces) and experimental observation (dashed traces and symbols) compared. Levelt I & II: weak increase of ⟨*T*⟩ with *c*_*dom*_ when *c*_*sup*_ =1.0 (red traces and symbols), and strong decrease with *c*_*sup*_ when *c*_*dom*_ =1.0 (brown traces and symbols). Levelt III: Symmetric increase of ⟨*T*⟩ with *c*_*dom*_ (red traces and symbols) and decrease with *c*_*sup*_ (brown traces and symbols), when *c*_*dom*_ +*c*_*sup*_ =100%. Alternation rate (green traces and symbols) peaks at equidominance and decreases symmetrically to either side. **(c)** Levelt IV: Decrease of ⟨*T*⟩ with image contrast, when *c*_*sup*_ =*c*_*sup*_. **(d)** Predicted dependence of sequential correlation *cc*_1_ (color scale) on *c*_*dom*_ and *c*_*sup*_. **(e)** Model prediction (black trace, *N* =25) and experimental observation (blue trace and symbols, mean±SEM, Spearman’s rank correlation *ρ*), when *c*_*sup*_ =*c*_*sup*_. Also shown is a second model prediction (red trace, *N* =40).

**Figure 3.**
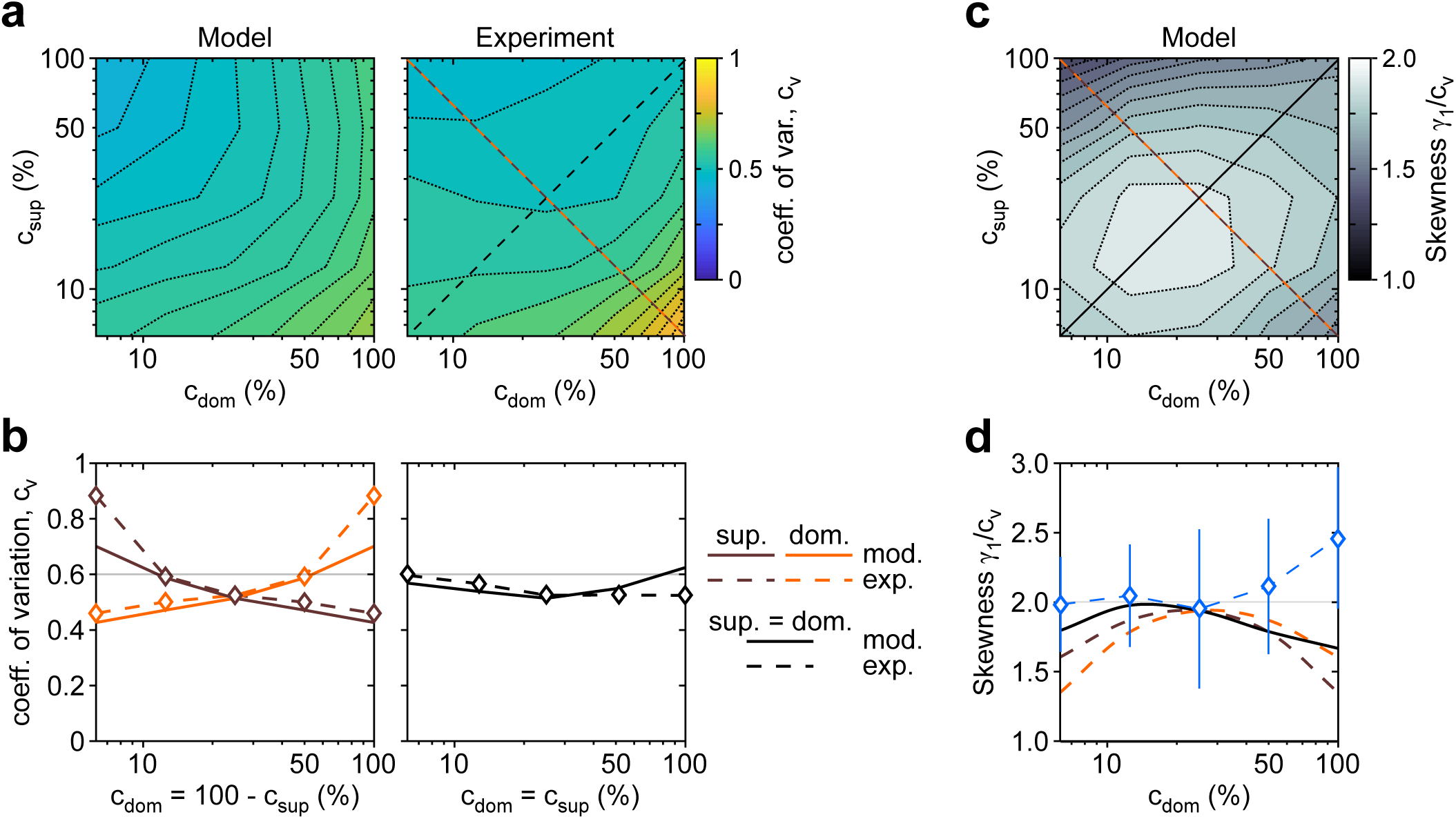
Shape of dominance distribution depends only weakly on image contrast (‘scaling property’). Distribution shape is parametrized by coefficient of variation *c*_*v*_ and relative skewness *γ*_1_*/c*_*v*_. **(a) (a)** Coefficient of variation *c*_*v*_ (color scale), as a function of dominant contrast *c*_*dom*_ and suppressed contrast *c*_*sup*_, in model (left) and experiment (right). **(b)** Model prediction (solid traces) and experimental observation (dashed traces and symbols) compared. Left: increase of *c*_*v*_ with *c*_*dom*_ (red traces and symbols), and symmetric decrease with *c*_*sup*_ (brown traces and symbols), when *c*_*sup*_ +*c*_*sup*_ =1.0. Right: weak dependence when *c*_*dom*_ =*c*_*sup*_ (black traces and symbols). **(c)** Predicted dependence of relative skewness *γ*_1_*/c*_*v*_ (gray scale) on *c*_*dom*_ and *c*_*sup*_. **(d)** Model prediction (solid traces) and experimental observation (blue dashed trace and symbols, mean±SEM), when *c*_*dom*_ = 1 − *c*_*sup*_.

**Figure 4.**
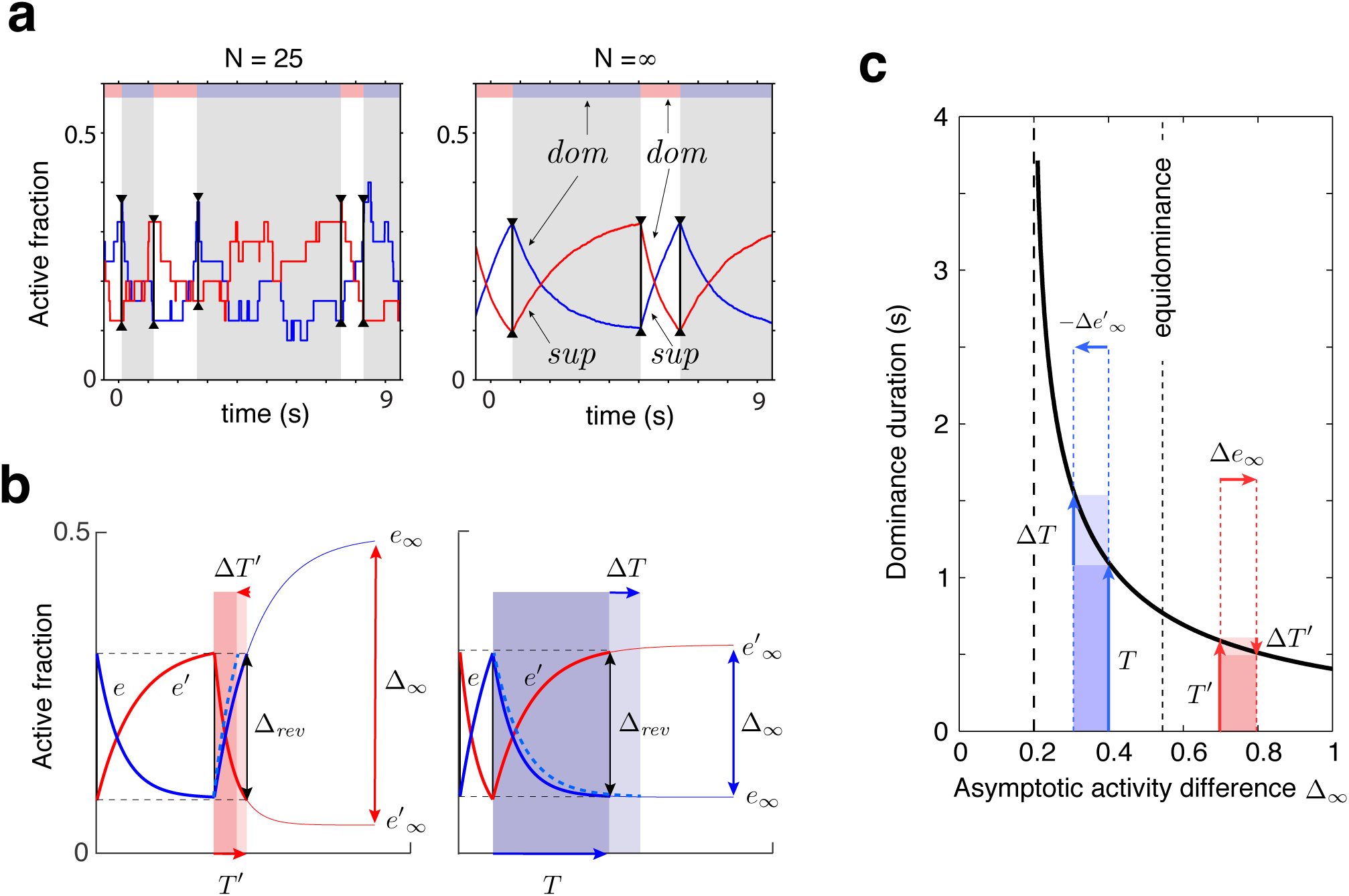
Joint dynamics of evidence habituation and recovery. Exponential development of evidence activities is governed by input-dependent asymptotic values and characteristic times. **(a)** Fractional activities *e* (blue traces) and *e*′ (red traces) of evidence pools *E* and *E*′, respectively, over several dominance periods for unequal stimulus contrast (*c*= 7*/*8, *c*′= 1*/*8). Stochastic reversals of finite system (*N* = 25 units per pool, left) and deterministic reversals of infinite system (*N* →∞, right). Perceptual dominance (decision activity) is indicated along the upper margin (red or blue stripe). Dominance evidence habituates (*dom*), and non-dominant evidence recovers (*sup*), until evidence contradicts perception sufficiently (black vertical lines) to trigger a reversal (grey and white regions). **(b)** Development of stronger-input evidence *e* (blue) and weaker-input evidence *e*′ (red), over two successive dominance periods (*c*= 15*/*16, *c*′= 1*/*16). Activities recover, or habituate, exponentially until reversal threshold Δ_*rev*_ is reached. Thin curves extrapolate to the respective asymptotic values, *e*_∞_ and *e*′_∞_. Dominance durations depend on distance Δ_∞_ and on characteristic times *τ*_*e*_ and *τ*_*e*′_. Left: incrementing non-dominant evidence *e* (dashed curve), raises upper asymptotic value *e*_∞_ and shortens dominance *T* ′ by Δ*T* ′. Right: incrementing dominant evidence *e* (dashed curve), raises lower asymptotic value *e*_∞_ and shortens dominance *T* by Δ*T*. **(c)** Increasing asymptotic activity difference Δ_∞_ accelerates the development of differential activity and curtails dominance periods *T, T* ′ (and *vice versa*). As the dependence is hyperbolic, any change to Δ_∞_ disproportionately affects longer dominance periods. If *T* >*T* ′, then Δ*T* >Δ*T* ′ (and *vice versa*).

**Figure 5.**
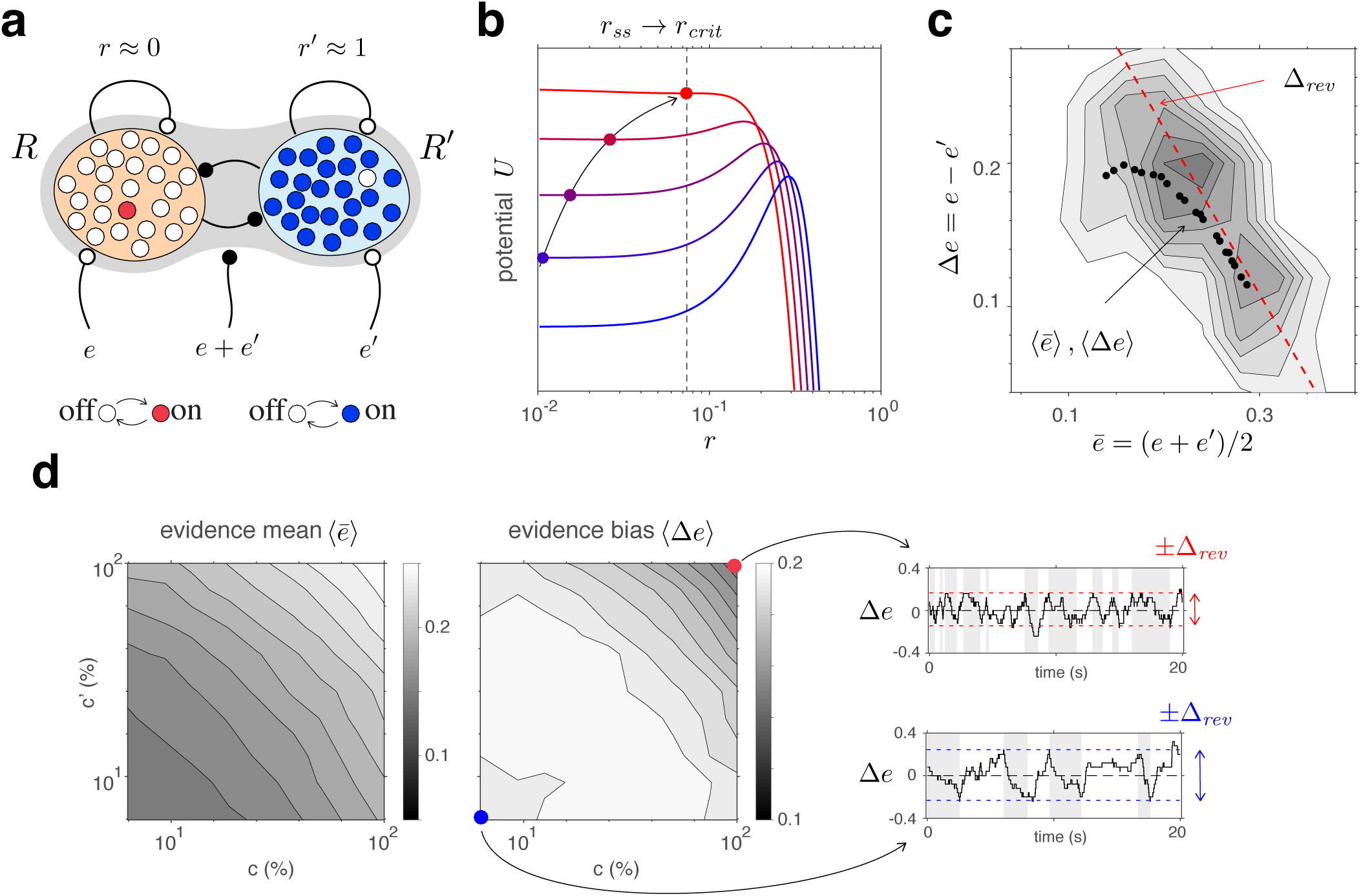
Competitive dynamics of decision pools ensures Levelt IV. **(a)** The joint stable state of decision pools (here *r*′ ≃ 1 and *r* ≃ 0) can be destabilized by sufficiently contradictory evidence, *e*>*e*′. **(b)** Effective potential *U* (*e, e*′, *r, r*′) (colored curves) and steady-states *r*_∞_ (colored dots) for different levels of contradictory input, Δ*e*=*e*−*e*′. Increasing Δ*e* destabilizes the steady-state and shifts *r*_∞_ rightwards (curved arrow). The critical value *r*_*crit*_ (dotted vertical line), at which the steady-state turns unstable, is reached when Δ*e* reaches the reversal threshold Δ_*rev*_. At this point, a reversal ensues with *r*→1 and *r*′→0. **(c)** The reversal threshold Δ_*rev*_ diminishes with combined evidence *e*+*e*′. In the deterministic limit, Δ_*rev*_ decreases linearly with 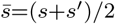 (dashed red line). In the stochastic system, the average evidence bias ⟨Δ*e*⟩ at the time of reversals decreases similarly with the average evidence mean *ē* (black dots). Actual values of Δ*e* at the time of reversals are distributed around these average values (grey shading). **(d)** Average evidence mean 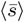 (left) and average evidence bias ⟨Δ*s*⟩ (middle) at the time of reversals, as a function of image contrast *c* or *c*′. Decrease of average evidence bias ⟨Δ*s*⟩ with contrast shortens dominance durations (Levelt IV). At low contrast (blue dot), higher reversal thresholds Δ_*rev*_ result in less frequent reversals (bottom right, grey and white regions) whereas, at high contrast (red dot), lower reversal thresholds lead to more frequent reversals (top right).

**Figure 6.**
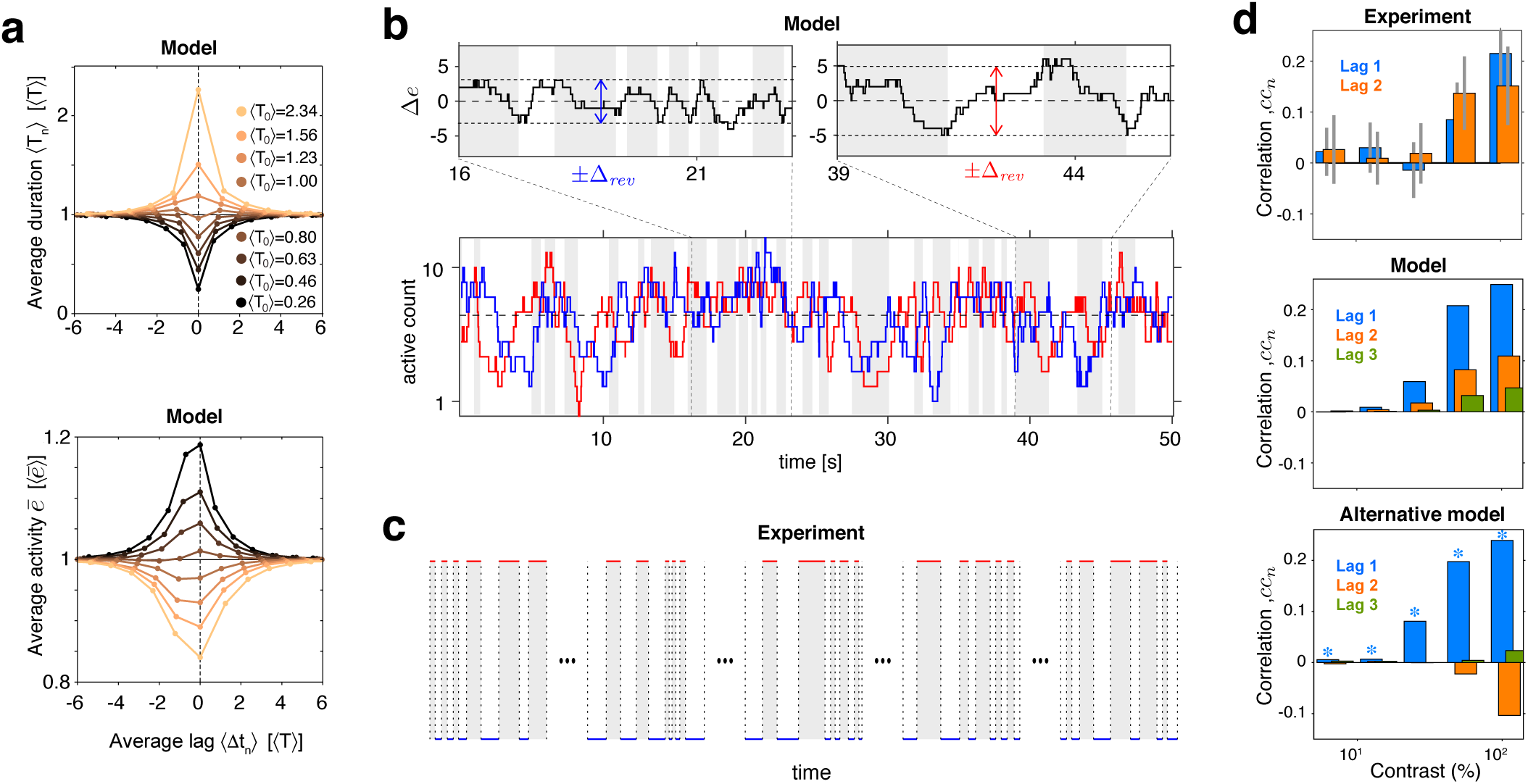
Serial dependency predicted by model and confirmed by experimental observations. **(a)** Conditional expectation of dominance duration ⟨*T*_±*n*_⟩ (top) and of mean evidence activity, 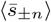 (bottom), in model simulations with maximal stimulus contrast (*c*=*c*′=1.0). Dominance periods *T*_0_ were grouped into octiles, from longest (yellow) to shortest (black). For each octile, the average duration ⟨*T*_±*n*_⟩ of preceding and following dominance periods, as well as the average mean evidence activity 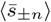 at the end of each period, are shown. All times in multiples of the overall average duration, ⟨*T*⟩. **(b)** Example reversal sequence from model. Bottom: Stochastic development of evidence activities *e* and *e*′ (red and blue traces), with large, joint fluctuations raising or lowering mean activity *ē*=(*e* + *e*′)*/*2 above or below long-term average (dashed line). Top left: episode with 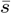 *above* average, *lower* Δ_*rev*_ and *shorter* dominance periods. Top right: episode with 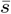 *below* average, *higher* Δ_*rev*_ and *longer* dominance durations. **(c)** Examples of reversal sequences from human observers (*c*=*c*′=1.0 and *c*=*c*′=0.5). **(d)** Positive lagged correlations predicted by model (mean, middle) and confirmed by experimental observations (mean±std, top). Alternative model [59] with adaptation & noise (mean, bottom), fitted separately at each contrast to reproduce ⟨*T*⟩, *c*_*v*_, *γ*_1_, and *cc*_1_ correlations of the present model (blue stars).

**Figure 7.**
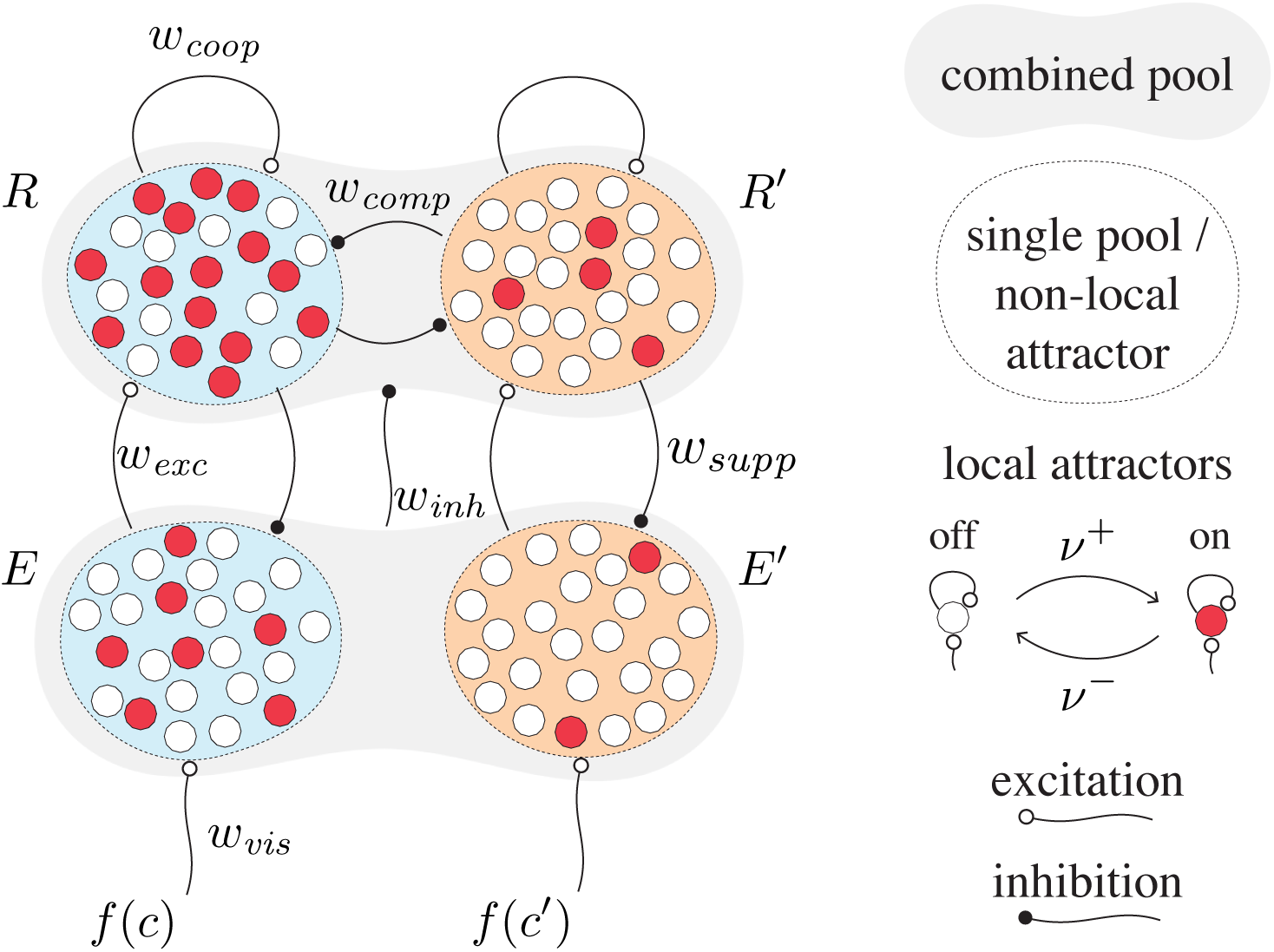
Proposed mechanism of binocular rivalry dyanmics (schematic). Bistable variables are represented by white (inactive) or red (active) circles. Four pools, each with *N* = 25 variables, are shown: two evidence pools *E* and *E*′, with active counts *n*_*e*_(*t*) and *n*_*e*′_ (*t*), and two decision pools, *R* and *R*′, with active counts *n*_*r*_(*t*) and *n*_*r*′_ (*t*). Excitatory and inhibitory synaptic couplings include selective feedforward excitation *w*_*exc*_, indiscriminate feedforward inhibition *w*_*inh*_, recurrent excitation *w*_*coop*_ and mutual inhibition *w*_*comp*_ of decision pools, as well as selective feedback suppression *w*_*supp*_ of evidence pools. Visual input to evidence pools *f* (*c*) and *f* (*c*′) is a function of image contrast *c* and *c*′.

**Figure 8.**
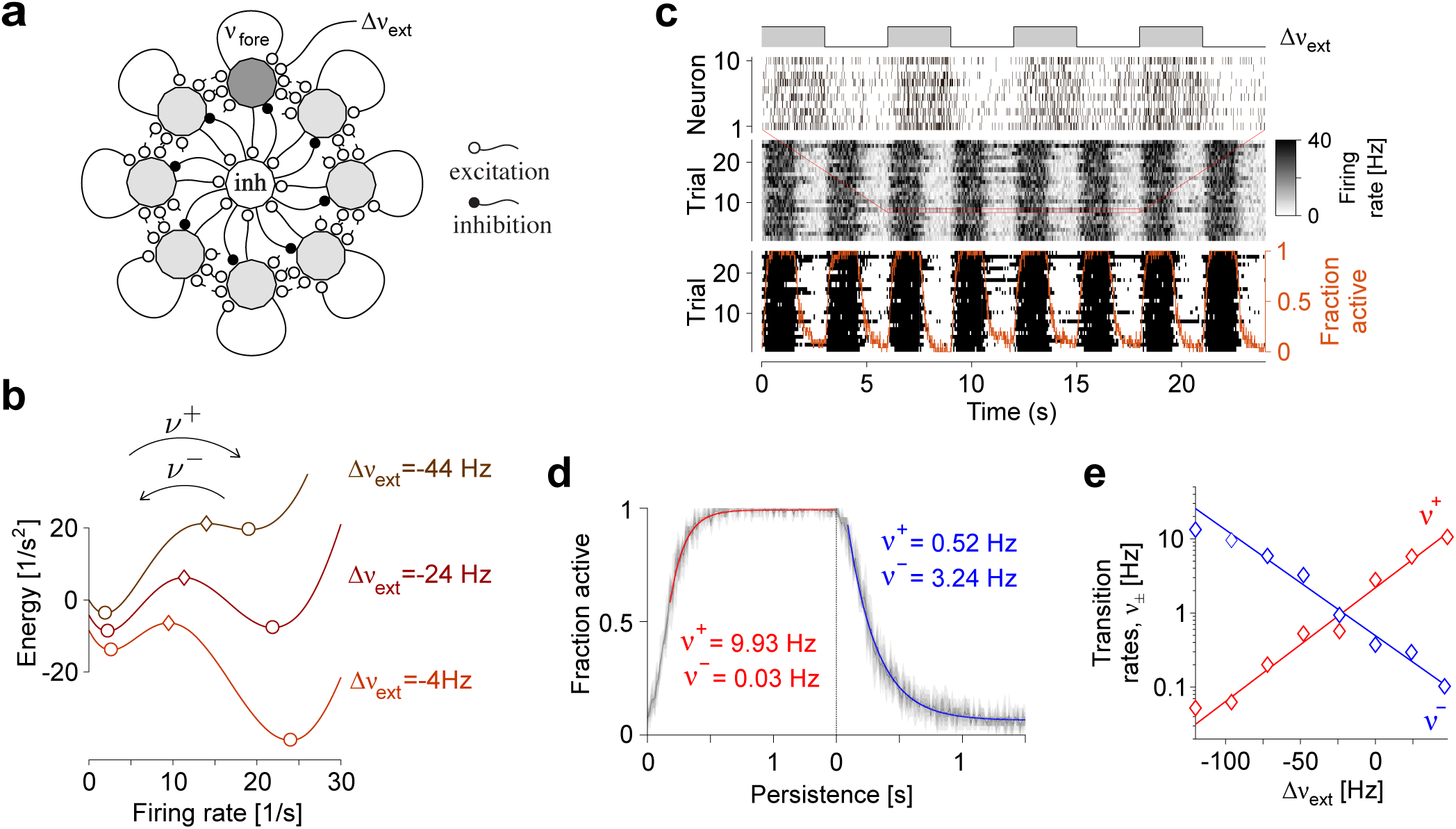
Metastable dynamics of spiking neural network. **(a)** Eight assemblies of excitatory neurons (schematic, light and dark gray disks) and one pool of inhibitory neurons (white disc) interact competitively with recurrent random connectivity. We focus on one ‘foreground’ assembly (dark gray), with firing rate *ν*_*fore*_ and selective external input Δ*ν*_*ext*_. **(b)** ‘Foreground’ activity explores an effective energy landscape with two distinct steady-states (circles), separated by ridge points (diamonds). As this landscape changes with external input Δ*ν*_*ext*_, transition rates *ν*_±_ between ‘on’ and ‘off’ states also change with external input. **(c)** Simulation to establish transition rates *ν*_±_ of fore-ground assembly. External input Δ*ν*_*ext*_ is stepped periodically between −44 *Hz* and −4 *Hz*. Spiking activity of 10 representative excitatory neurons in a single trial, population activity over 25 trials, thresholded population activity over 25 trials, and activation probability (fraction of ‘on’ states). **(d)** Relaxation dynamics in response to step change of Δ*ν*_*ext*_, with ‘on’ transitions (left) and ‘off’ transitions (right). **(e)** Average state transition rates *ν*^±^ vary exponentially and symmetrically with external input. The dependence is well approximated by *ν*^±^≃ 1*Hz* exp (±0.04*s*Δ*ν*_*ext*_ + 1) (red and blue lines).

#### Binary stochastic variables

Our model reduces bistable assemblies to discretely stochastic, binary activity variables *x*(*t*)∈{0, 1}, which activate and inactivate with Poisson rates rates *ν*^+^ and *ν*^−^, respectively. These rates *ν*^±^(*s*) vary exponentially with activation energy Δ*u*=*u*(*s*) + *u*_0_:

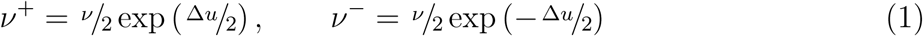

where *u*_0_ and *ν* are baseline potential and and baseline rate, respectively, and where input-dependent part *u*(*s*)=*ws* varies linearly with input *s*, with synaptic coupling constant *w* (supplementary section **Metastable population dynamics** and **Fig. 8e**).

#### Pool of N binary variables

An extended network, containing *N* individually bistable assemblies with shared input *s*, reduces to a ‘pool’ of *N* binary activity variables *x*_*i*_(*t*)∈{0, 1} with identical rates *ν*_±_(*s*). Although all variables are independently stochastic, they are coupled through their shared input *s*. The number of active variables *n*(*t*)= Σ*i x*_*i*_(*t*) or, equivalently, the active fraction *x*(*t*)=*n*(*t*)*/N*, forms a discretely stochastic process (“birth-death” or “Ehrenfest” process [49]).

#### Relaxation dynamics

While activity *x*(*t*) develops discretely and stochastically according to Eq. 5 (Methods), its expectation ⟨*x*(*t*)⟩ develops continuously and deterministically,

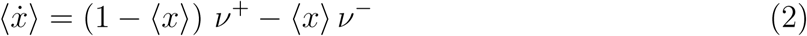

relaxing with characteristic time 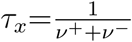 towards asymptotic value 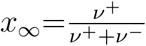. As rates *ν*^±^ change with input *s* (Eq. 1), so do *τ*_*x*_=ϒ (*s*) and *x*_∞_=Φ(*s*) (see Methods). Characteristic time *τ*_*x*_ is longest for small input *s*≃0 and shortens for larger positive or negative input |*s*|>>0. Asymptotic value *x*_∞_ ranges over the interval [0, 1] and varies sigmoidally with input *s*, reaching half-activation for *s*=−*u*_0_*/w*.

#### Quality of representation

Pools of bistable variables belong to a class of neural representations particularly suited for Bayesian integration of sensory information [6, 93]. In general, summation of activity is equivalent to optimal integration of information, provided that response variability is Poisson-like, and response tuning differs only multiplicatively [70, 71]. Pools of bistable variables closely approximate these properties (see supplementary text **Quality of representation:** *Suitability for inference*).

The representational accuracy of even a comparatively small number of bistable variables can be surprisingly high. For example, if normally distributed inputs drive the activity of initially inactive pools of bistable variables, pools as used in the present model (*N* =25, *w*=2.5) readily capture 90% of the Fisher information (supplementary text **Quality of representation:** *Integration of noisy samples*).

#### Conflicting evidence

Any model of binocular rivalry must represent the conflicting evidence from both eyes (*e.g*., different visual orientations), which supports alternative perceptual hypotheses (*e.g*., distinct grating patterns). We assume that conflicting evidence accumulates in two separate pools of *N* =25 bistable variables, *E* and *E*′, (‘evidence pools’, **Fig. 1c**). Fractional activations *e*(*t*) and *e*′(*t*) develop stochastically following Eq. 5 (Methods). Transition rates 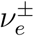 and 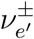 vary exponentially with activation energy (Eq. 1), with baseline potential 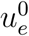 and baseline rate *ν*_*e*_. The variable components of activation energy, *u*_*e*_ and *u*_*e*′_, are synaptically modulated by image contrasts, *c* and *c*′:

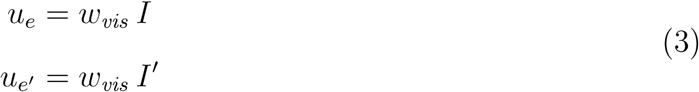

where *w*_*vis*_ is a coupling constant and *I*=*f* (*c*)∈[0, 1] is a monotonic function of image contrast *c*∈[0, 1] (see Methods).

#### Competing hypotheses: ‘non-local attractors’

Once evidence for, and against, alternative perceptual hypotheses (*e.g*., distinct grating patterns) has been accumulated, reaching a decision requires a sensitive and reliable mechanism for identifying the best supported hypothesis and amplifying the result into a categorical read-out. Such a winner-take-all decision [52] is readily accomplished by a dynamical version of biased competition [30, 31, 120, 121].

We assume that alternative perceptual hypotheses are represented by two further pools of *N* = 25 bistable variables, *R* and *R*′, forming two ‘non-local attractors’ (‘decision pools’, **Fig. 1c**). Similar to previous models of decision making and attentional selection [30, 31, 120, 121], we postulate recurrent excitation within pools, but recurrent inhibition between pools, to obtain a ‘winner-take-all’ dynamics. Importantly, we assume that ‘evidence pools’ project to ‘decision pools’ not only in the form of selective excitation (targeted at the corresponding decision pool), but also in the form of indiscriminate inhibition (targeting both decision pools), as suggested previously [10, 34].

Specifically, fractional activations *r*(*t*) and *r*′(*t*) develop stochastically according to Eq. 5 (Methods). Transition rates 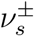 and 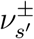 vary exponentially with activation energy (Eq. 1), with baseline difference 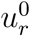 and baseline rate *ν*_*r*_. The variable components of activation energy, *u*_*r*_ and *u*_*r*′_, are synaptically modulated by evidence and decision activities:

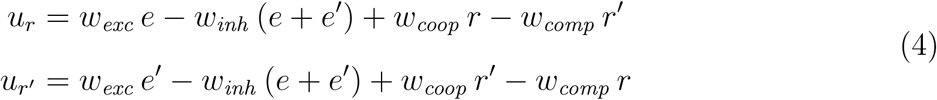

where coupling constants *w*_*exc*_, *w*_*inh*_, *w*_*coop*_, *w*_*comp*_ reflect feedforward excitation, feedforward inhibition, lateral cooperation within decision pools, and lateral competition between decision pools, respectively.

This biased competition circuit expresses a categorical decision by either raising *r* towards unity (and lowering *r*′ towards zero) or *vice versa*. The choice is random when visual input is ambiguous, *I*≃*I*′, but becomes deterministic with growing input bias |*I* −*I*′|>0. This probabilistic sensitivity to input bias is reliable and robust under arbitrary initial conditions of *e, e*′, *r* and *r*′ (see supplementary section **Categorical choice** with **Fig. 10**).

**Figure 9.**
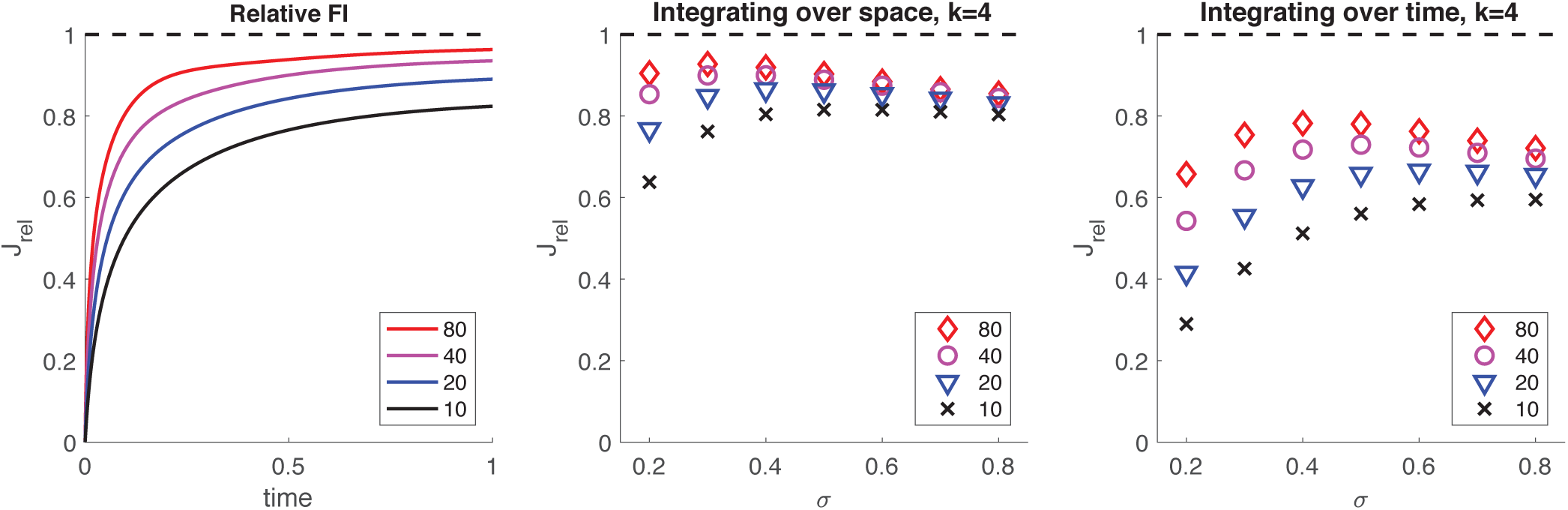
Supplementary. Information retained by stochastic pool activity from normally distributed inputs. Inputs *s* ∈ 𝒩 (*µ, σ*) provide Fisher information *J*_*s*_ = 1*/σ*^2^ about mean *µ*. Stochastic activity *n*(*t*) of a birth-death process (*N* ∈ {10, 20, 40, 80} and *w* = 2.5) driven by such inputs accumulates Fisher information *J*_*n*_(*t*) about mean *µ*. **(a)** Accumulation over input interval *t* = [0, 1] of fractional information *J*_*rel*_ (*t*) = *J*_*n*_(*t*) *σ*^2^ by an initially inactive pool of size *N*. **(b)** Information about *µ* retained by *summed activity* 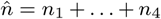 of four independent pools (all initially inactive and of size *N*) receiving concurrently four independent inputs (*s* ∈ *𝒩* (*µ, σ*)) over an interval *t* = [0, 1]. Retained fraction 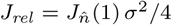 (1) *σ*^2^*/*4 depends on pool size *N* and input variance *σ*^2^. **(c)** Information about *µ* retained by activity *n* of one pool (initially inactive and of size *N*) receiving successively four independent inputs (*s* ∈ 𝒩 (*µ, σ*)) over an interval *t* = [0, 43]. Retained fraction *J*_*rel*_ = *J*_*n*_(4) *σ*^2^*/*4 depends on pool size *N* and input variance *σ*^2^.

**Figure 10.**
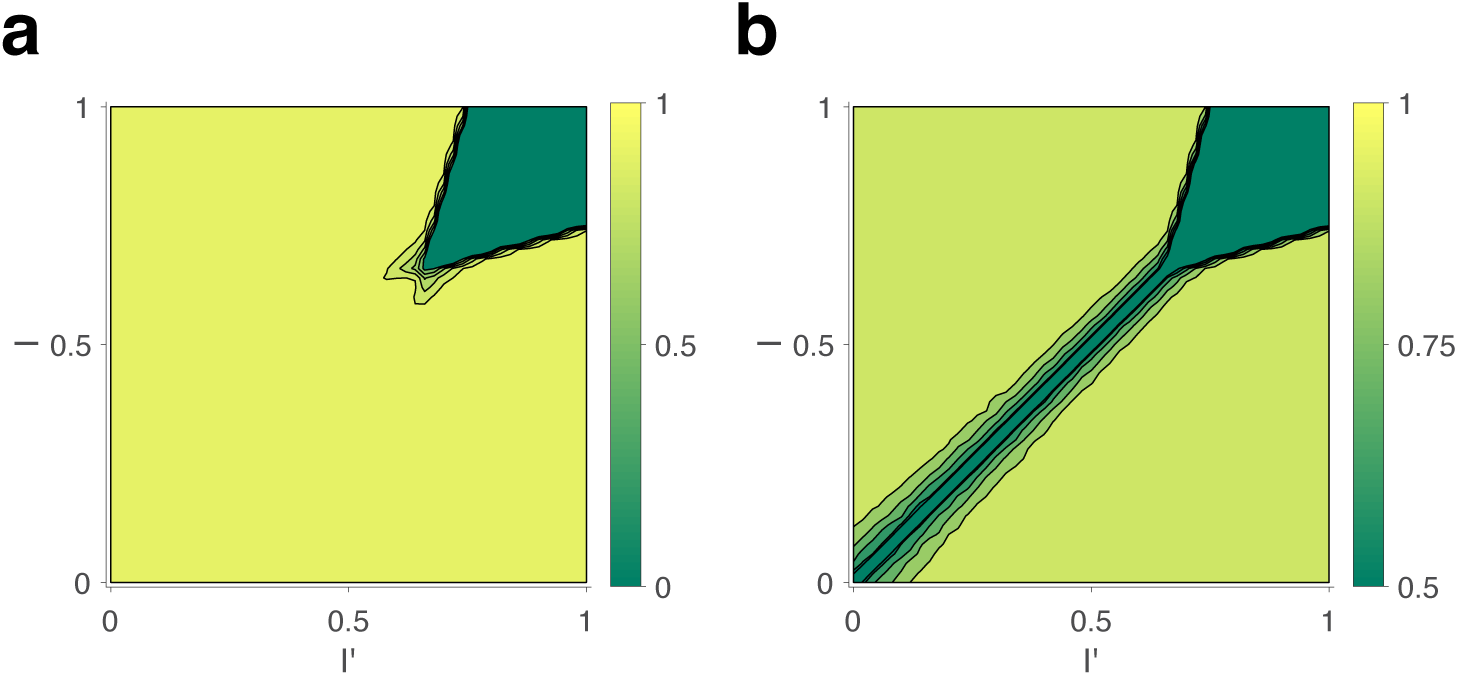
Decision response to fixed input *I, I*′, for random initial conditions of *e, e*′, *r, r*′. (a) Expected differential steady-state activation ⟨|*r* − *r*′|⟩ of decision level. Steady-state activity *r* + *r*′ ≃ 1 implies a categorical decision with activity ≃ 1 of one pool and activity ≃ 0 of another. (b) Probability that decision correctly reflects input bias (*r* > *r*′ if *I* > *I*′, and *vice versa*).

#### Feedback suppression

Finally, we assume feedback suppression, with each decision pool selectively targeting the corresponding evidence pool. A functional motivation for this systematic bias *against* the currently dominant appearance is given momentarily. Its effects include curtailment of dominance durations and ensuring that reversals occur from time to time. Specifically, we modify Eq. 3 to

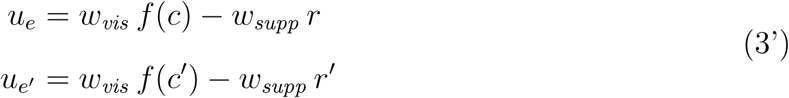

where *w*_*supp*_ is a coupling constant.

Previous models of binocular rivalry [27, 46] have justified selective feedback suppression of the evidence supporting a winning hypothesis in terms of “predictive coding” and “hierarchical Bayesian inference” [61, 97]. An alternative normative justification is that, in volatile environments, where the sensory situation changes frequently (‘volatility prior’), optimal inference requires an exponentially growing bias *against* evidence for the most likely hypothesis [117]. Note that feedback suppression applies selectively to evidence for a winning hypothesis and is thus materially different from visual adaptation [123], which applies indiscriminately to all evidence present.

### Reversal dynamics

A representative example of the joint dynamics of evidence and decision pools is illustrated in **Fig. 1cd**, both at the level of pool activities *e*(*t*), *e*′(*t*), *r*(*t*), *r*′(*t*), and at the level of individual bistable variables *x*(*t*). The top row shows decision pools *R* and *R*′, with instantaneous active counts, *Nr*(*t*) and *Nr*′(*t*) and active / inactive states of individual variables *x*(*t*). The bottom row shows evidence pools *E* and *E*′, with instantaneous active counts, *Ne*(*t*) and *Ne*′(*t*) and active / inactive states of individual variables *x*(*t*). Only a small fraction of evidence variables is active at any one time.

Phenomenal appearance reverses when the differential activity Δ*e*=*e*-*e*′ of evidence pools, *E* and *E*′, *contradicts* sufficiently strongly the differential activity Δ*r*=*r*-*r*′ of decision pools, *R* and *R*′, such that the steady-state of decision pools is destabilized (see further below and **Fig. 5**). As soon as the reversal has been effected at the decision level, feedback suppression *lifts from* the newly non-dominant evidence, and *descends upon* the newly dominant evidence. Due to this asymmetric suppression, the newly non-dominant evidence recovers, whereas the newly dominant evidence habituates. This opponent dynamics progresses, past the point of equality *s*∼*s*′, until differential evidence activity Δ*e*=*e*-*e*′ once again *contradicts* differential decision activity Δ*r*=*r*-*r*′. Accordingly, the respective times at which evidence and decision activities reverse are phase-shifted by approximately 90° relative to each other (**Fig. 1d**), consistent with previous models [1, 41, 124].

### Binocular rivalry

To compare predictions of the model described above to experimental observations, we measured spontaneous reversals of binocular rivalry for different combinations of image contrast. Binocular rivalry is a particularly well studied instance of multistable perception [15, 33, 65, 66, 126]. When conflicting images are presented to each eye (*e.g*., by means of a mirror stereoscope or of colored glasses, see Methods), the phenomenal appearance reverses from time to time between the two images (**Fig. 1a**). Importantly, the perceptual conflict involves also representations of coherent (binocular) patterns and is not restricted to eye-specific (monocular) representations [7, 11, 54, 68].

Specifically, our experimental observations established reversal sequences for 5 × 5 combinations of image contrast, *c*_*dom*_, *c*_*sup*_ ∈ { 1*/*16, 1*/*8, 1*/*4, 1*/*2, 1}, where *c*_*dom*_ pertains to the phenomenally dominant image and *c*_*sup*_ to the phenomenally suppressed image (see Methods). We analysed these observations in terms of mean dominance durations ⟨*T*⟩, higher moments *c*_*v*_ and *γ*_1_*/c*_*v*_ of the distribution of dominance durations, and sequential correlation *cc*_1_ of successive dominance durations. Additional aspects of serial dependence are discussed further below.

As described in Methods, we fitted twelve (12) model parameters to reproduce observations with more than 50 degrees of freedom: 5 × 5 mean dominance durations ⟨*T*⟩, coefficients of variation *c*_*v*_, and values of skewness *γ*_1_*/c*_*v*_=2, as well as a uniform correlation coefficient *cc*_1_=0.06. Model predictions and experimental observations are juxtaposed in **Fig. 2** and **Fig. 3**.

The complex and asymmetric dependence of mean dominance durations on image contrast — aptly summarized by Levelt’s ‘propositions’ I to IV [15, 66] — is fully reproduced by the model (**Fig. 2a**). Increasing the contrast of one image, increases the fraction of time during which this image dominates appearance (‘predominance’, Levelt I). Counter-intuitively, this is due more to shortening dominance of the *unchanged* image than to lengthening dominance of the *changed* image (Levelt II, **Fig. 2b**, left panel). Mean dominance durations grow (and alternation rates decline) symmetrically around equal predominance as contrast difference *c*_*dom*_ −*c*_*sup*_ increases (Levelt III, **Fig. 2b**, right panel). Mean dominance durations shorten when both image contrasts *c*_*dom*_ =*c*_*sup*_ increase (Levelt IV, **Fig. 2c**).

Successive dominance dominance durations are typically correlated positively [39, 89, 119]. Averaging over all contrast combinations, observed and predicted correlation coefficients were comparable with *cc*_1_ = 0.06±0.06 (mean and standard deviation). Unexpectedly, both observed and predicted correlations coefficients increased systematically with image contrast (*ρ* = 0.9, *p* < .01), growing from *cc*_1_ = 0.02 ± 0.05 at *c*_*dom*_ =*c*_*sup*_=1*/*_16_ to 0.21 ± 0.06 at *c*_*dom*_ =*c*_*sup*_=1 (**Fig. 2e**, blue symbols). To our knowledge, this contrast-sensitivity of sequential correlations has not been observed previously.

The distribution of dominance durations typically takes a characteristic shape [8, 12, 13, 19, 28, 32, 39, 82, 88, 119], approximating a Gamma distribution with shape parameter *r* ≃ 3−4, or coefficient of variation 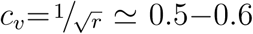. The fitted model fully reproduces this ‘scaling property’ (**Fig. 3a**). The observed coefficient of variation remained in the range *c*_*v*_ ≃ 0.5−0.6 for nearly all contrast combinations (**Fig. 3b**). Unexpectedly, both observed and predicted values increased above, or decreased below, this range at extreme contrast combinations (**Fig. 3b**, left panel). At intermediate contrast, where observed values of skewness were most reliable, both observed and predicted values of skewness were *γ*_1_*/c*_*v*_≃2 and conformed to a Gamma distribution (**Fig. 3d**, blue symbols).

### Reasons for success

What are the reasons for the surprising success of the model described above? Which features of its dynamics are responsible for reproducing different aspects of binocular rivalry? Here we explain why no component of the model may reasonably be omitted and how highly non-trivial interactions between these components guarantee the universal characteristics of multistable phenomena.

#### Levelt’s propositions I, II and III

The characteristic input-dependence of average dominance durations emerges in two steps. First, inputs and feedback suppression shape the birth-death dynamics of evidence pools *E* and *E*′ (by setting disparate transition rates *ν*^±^, following Eq. 3’ and Eq. 1). Second, this sets in motion two opponent developments (habituation of dominant evidence activity and recovery of non-dominant evidence activity, both following Eq. 2) that jointly determine dominance duration.

To elucidate this mechanism, it is helpful to consider the limit of large pools (*N* → ∞) and its deterministic dynamics (**Fig. 4a**), which corresponds to the *average* stochastic dynamics. In this limit, periods of dominant evidence *E* or *E*′ start and end at the same levels (*e*_*start*_ =*e*′_*start*_ and *e*_*end*_ =*e*′_*end*_), because reversal thresholds Δ_*rev*_ are the same for evidence difference *e*−*e*′ and *e*′−*e* (see section *Levelt IV* below).

The rates at which evidence habituates or recovers depend, in the first instance, on asymptotic levels *e*_∞_ and *e*′_∞_ (Eq. 1 and 2, **Fig. 4b** and **Fig. 11**). In general, dominance durations depend on distance Δ_∞_ between asymptotic levels: the further apart these are, the faster the development and the shorter the duration. As feedback suppression *inverts* the sign of the opponent developments, dominant evidence decreases (habituates) while nondominant evidence increases (recovers). Due to this inversion, Δ_∞_ is roughly proportional to 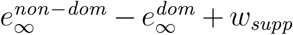. It follows that distance Δ_∞_ is *smaller* and the reversal dynamics *slower* when dominant input is *stronger*, and *vice versa*. It further follows that incrementing one input (and raising the corresponding asymptotic level), speeds up recovery or slows down habituation, shortening or lengthening periods of non-dominance and dominance, respectively (Levelt I).

**Figure 11.**
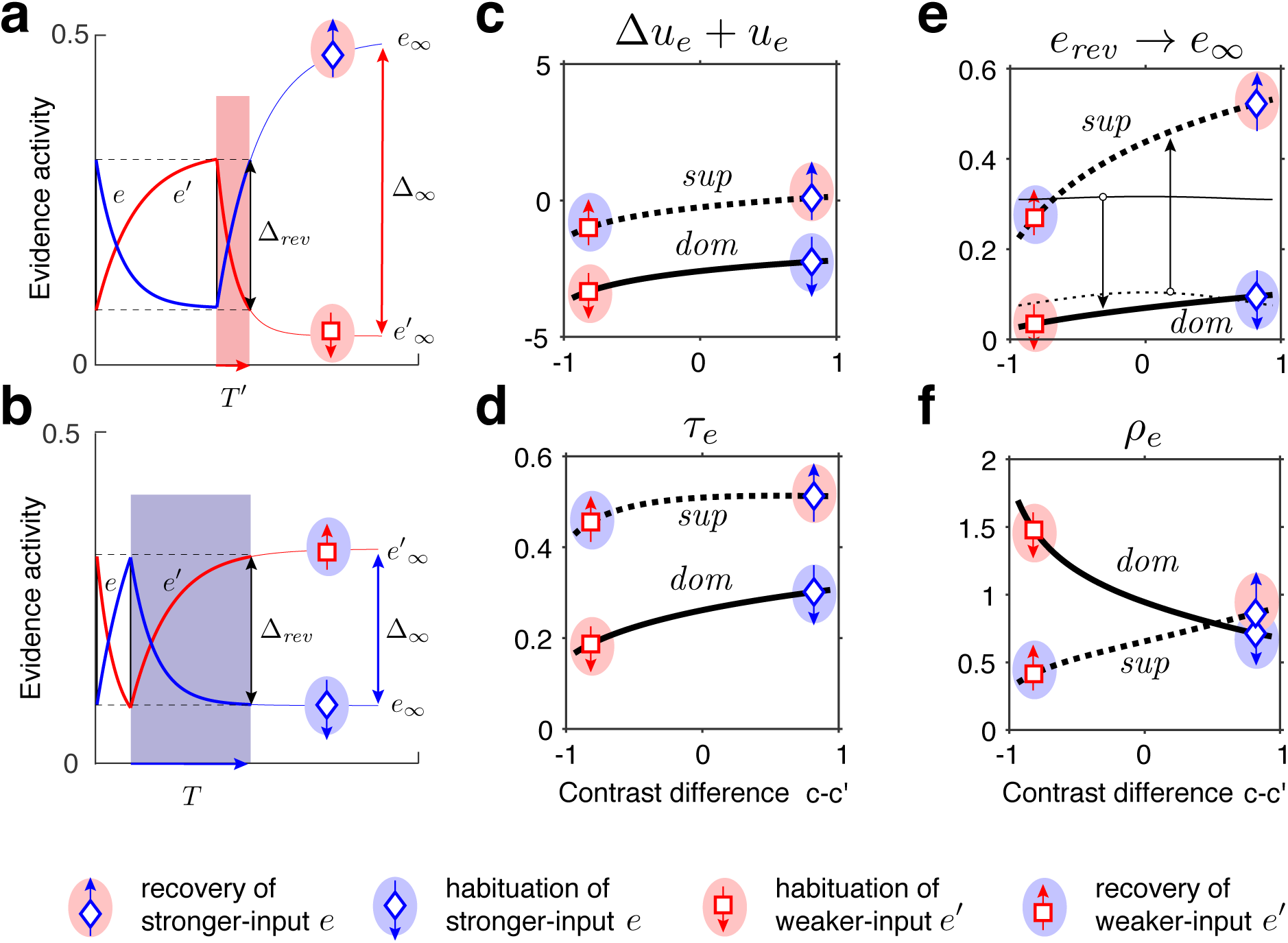
Exponential habituation and recovery of evidence activities. Dominance durations depend on distance between asymptotic values and on characteristic times. **(ab)** Development of evidence *e* (blue) and *e*′ (red), over two successive dominance periods. Input *c*=0.9375 is stronger, input *c*′ = 0.0625 weaker. Activities recover, or habituate, exponentially until reversal threshold Δ_*rev*_ is reached. Thin curves extrapolate to the respective asymptotic values, *e*_∞_ and *e*′_∞_. **(a)** Evidence *e*′ (with weaker input *c*′) is dominant. Incrementing input *c* to non-dominant evidence *e* shortens dominance *T* ′. **(b)** Evidence *e* (with stronger input *c*) is dominant. Incrementing input *c* to *e* extends dominance *T*. **(c-f)** Contrast-dependence of relaxation dynamics, as a function of differential contrast *c*−*c*′, for *c*+*c*′=1. Values when evidence *e* is dominant (*dom*, thick solid curves), and when it is non-dominant (*sup*, thick dotted curves). Values for *e*′ are mirror symmetric (about vertical midline *c*−*c*′=0). **(c)** Effective potential Δ*u*_*e*_ + *u*_*e*_. **(d)** Characteristic time *τ*_*e*_. **(e)** Relaxation range *e*_*rev*_ →*e*_∞_ (bottom left, thin curves *e*_*rev*_, thick curves *e*_∞_). **(f)** Effective rate *ρ*_*e*_ of development. Symbols and arrows correspond to subfigures **(ab)** and represent recovery (up arrow) or habituation (down arrow) of stronger-input evidence (blue) or weaker-input evidence (red). Underlying color patches indicate dominance of stronger-input evidence (blue patches) or of weaker-input evidence (red patches). Dominance durations depend more sensitively on the slower development, with smaller *ρ*, which generally is the recovery of non-dominant evidence (up arrows).

In the second instance, rates of habituation or recovery depend on characteristic times *τ*_*e*_ and *τ*_*e*′_ (Eq. 1 and 2). When these rates are unequal, dominance durations depend more sensitively on the *slower* process. This is why dominance durations depend more sensitively on non-dominant input (Levelt II): recovery of non-dominant evidence is generally slower than habituation of dominant evidence, independently of which input is weaker or stronger. The reason is that the respective effects of characteristic times *τ*_*e*_ and *τ*_*e*′_ and asymptotic levels *e*_∞_ and *e*′_∞_ are synergistic for weaker-input evidence (in both directions), whereas they are antagonistic for stronger-input evidence (supplementary text **Deterministic dynamics:** *Evidence pools* and **Fig. 11**).

In general, dominance durations depend hyperbolically on Δ_∞_ (**Fig. 4d** and Eq. 7 in supplementary text). Dominance durations become infinite (and reversals cease) when Δ_∞_ falls below the reversal threshold Δ_*rev*_. This hyperbolic dependence is also why alternation rate peaks at equidominance (Levelt III): increasing the difference between inputs always lengthens longer durations more than it shortens shorter durations, thus lowering alternation rate.

#### Distribution of dominance durations

For all combinations of image contrast, the mechanism accurately predicts the experimentally observed distributions of dominance durations. This is owed to the stochastic activity of pools of bistable variables.

Firstly, dominance distributions retain nearly the same shape, even though average durations vary more than three-fold with image contrast (supplementary **Fig. 12ab**). This ‘scaling property’ is due to the Poisson-like variability of birth-death processes (see supplementary text **Stochastic dynamics**). Generally, when a stochastic accumulation approaches threshold, the rates of both accumulation and dispersion of activity affect the distribution of first-passage-times [18, 19]. In the special case of Poisson-like variability, the two rates vary proportionally and preserve distribution shape (supplementary **Fig. 12cd**).

Secondly, predicted distributions correspond to Gamma distributions with scale factor *r*≃3-4. As shown previously [18, 19], this is due to birth-death processes accumulating activity within a narrow range (*i.e*., evidence difference Δ*e*≲0.2). In this low-threshold regime, the first-passage-times of birth-death processes are both highly variable and Gamma distributed, consistent with experimental observations.

Thirdly, the predicted variability (coefficients of variation) of dominance periods varies along the *c*+*c*′=1 axis, being larger for longer than for shorter dominance durations (**Fig. 3ab**). The reason is that stochastic development becomes noise-dominated. For longer durations, stronger-input evidence habituates rapidly into a regime where random fluctuations gain importance (see also supplementary **Fig. 11ab**).

#### Levelt’s proposition IV

The model accurately predicts how dominance durations shorten with higher image contrast *c*=*c*′ (Levelt IV). Surprisingly, this reflects the dynamics of decision pools *R* and *R*′ (**Fig. 5**).

Here again it is helpful to consider the deterministic limit of large pools (*N* →∞). In this limit, a dominant decision state *r*′≃1 is destabilized when a *contradictory* evidence difference Δ*e*=*e*−*e*′ exceeds a certain threshold value Δ_*rev*_ (**Fig. 5b** and supplementary text **Deterministic dynamics:** *Decision pools*). Due to the combined effect of excitatory and inhibitory feedforward projections, *w*_*exc*_ and *w*_*inh*_ (Eq. 4 and **Fig. 5a**), this average reversal threshold *decreases* with average evidence activity (*e*+*e*′)*/*2. Simulations of the fully stochastic model (*N* = 25) confirm this analysis (**Fig. 5c**). As average evidence activity (*e*+*e*′)*/*2 increases with image contrast, the average evidence bias Δ*e* at the time of reversals decreases, resulting in shorter dominance periods (**Fig. 5d**).

### Serial dependence

The proposed mechanism accurately predicts positive correlations between successive dominance durations, a well-known characteristic of multistable phenomena [32, 39, 114, 119]. In addition, it predicts further aspects of serial dependence not described previously.

In both model and experimental observations, a long dominance periods tends to be followed by another periods, and a short dominance periods by another short period (**Fig. 6**). In the model, this is due to mean evidence activity *ē*=(*e* + *e*′)*/*2 fluctuating stochastically above and below its long-term average. The autocorrelation time of these fluctuations increases monotonically with image contrast and, for high contrast, spans multiple dominance periods (supplementary text **Characteristic times** and **Fig. 13**). Note that fluctuations of *ē* diminish as the number of bistable variables increases and vanish in the deterministic limit (*N* → ∞).

**Figure 12.**
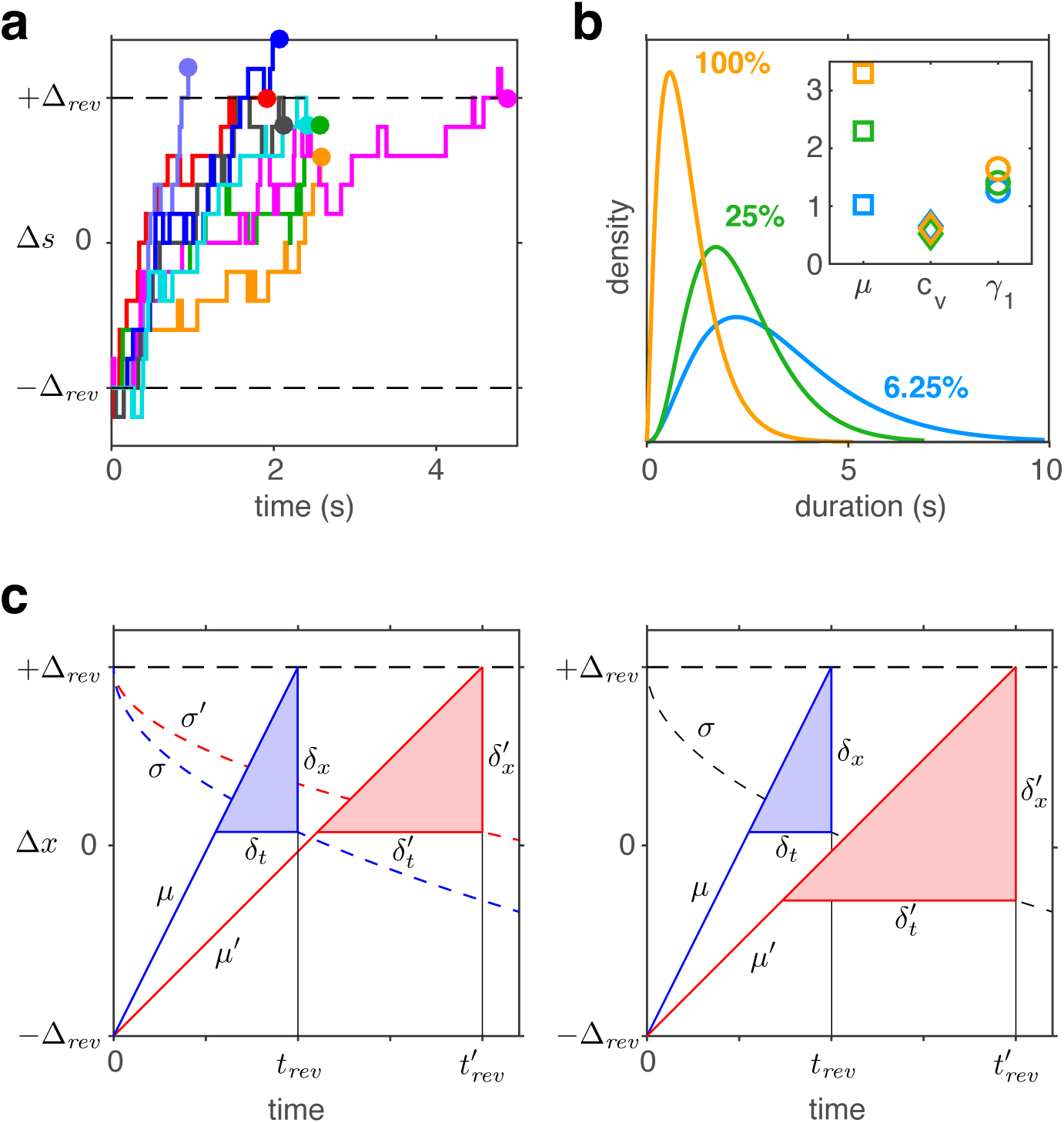
Birth-death dynamics of evidence pools ensures Gamma-like distribution and ‘scaling property’ (invariance of distribution shape). **(a)** Representative examples for the time-development of evidence bias Δ*e*=*e* − *e*′ between reversals (*i.e*., between −Δ_*rev*_ and approximately +Δ_*rev*_). (b) Dominance distributions for *c*=*c*′=0.0625 (blue), *c*=*c*′=0.25 (green), and *c*=*c*′=1.0 (yellow). Distribution mean *µ* changes approximately threefold, but coefficient of variation *c*_*v*_ and skewness *γ*_1_ are nearly invariant (inset), largely preserving distribution shape. **(c)** Development of expectation ⟨Δ*x*⟩ between reversals (schematic). Left: a Poisson-variable process, such as the difference Δ*x* between two birth-death processes. Mean ⟨Δ*x*⟩ grows linearly with *t* (lines, with slopes *µ, µ*′) and variance ⟨(Δ*x* − ⟨Δ*x*⟩)^2^⟩ grows linearly with 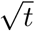 (dashed curves, with scaling factors *σ, σ*′). Constants *µ* and *σ* change with stimulus contrast (blue and red). Proportionality *µ* ∝ *σ*^2^, ensures constant dispersion of Δ*x* at threshold 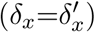, and, consequently, a dispersion of threshold-crossing times that grows linearly with mean threshold-crossing time 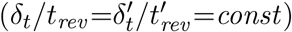, preserving distribution shape. Right: a process with constant variance, *σ*=*σ*′. Dispersion of Δ*x* at threshold increases with threshold-crossing time 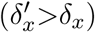 and dispersion of threshold-crossing times grows supra-linearly with mean threshold-crossing time 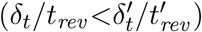, broadening distribution shape.

**Figure 13.**
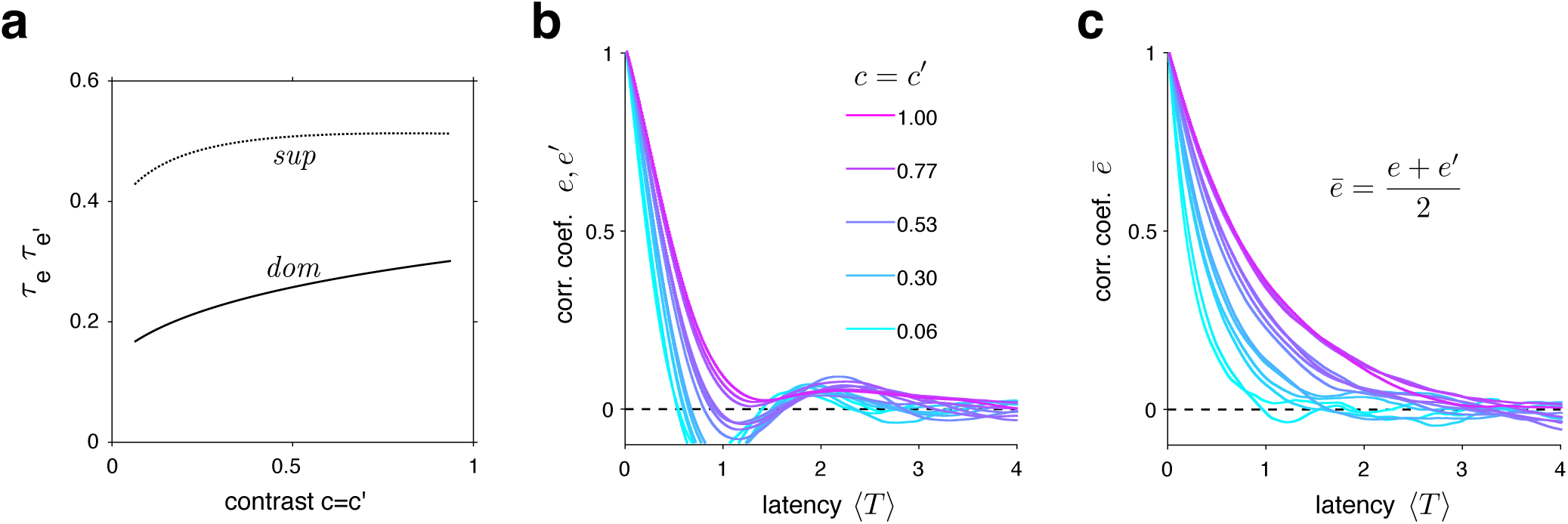
Characteristic times of evidence activity. **(a)** Characteristic times *τ*_*e*_, *τ*_*e*′_ for different image contrast *c*=*c*′, when evidence pool is dominant (*dom*) and non-dominant (*sup*). **(b)** Auto-correlation of evidence activity *e, e*′ as a function of image contrast (color) and latency, expressed in multiples of average dominance duration ⟨*T*⟩. **(b)** Autocorrelation of joint evidence activity *ē*=(*e* + *e*′)*/*2 as a function of image contrast (color) and latency. Note that autocorrelation time lengthens substantially for high image contrast.

Crucially, fluctuations of *ē* modulate both reversal threshold Δ_*rev*_ and dominance durations *T*, as illustrated in **Fig. 6ab**. To obtain **Fig. 6a**, dominance durations were grouped into quantiles and the average duration ⟨*T*_0_⟩ of each quantile was compared to the conditional expectation of preceding and following durations ⟨*T*_±*n*_⟩ (upper graph). For the same quantiles (compare color coding), average evidence activity *ē*_0_ was compared to the conditional expectation ⟨*ē*_±*n*_⟩ at the end of preceding and following periods (lower graph). Both the *inverse* relation between ⟨*T*_±*n*_⟩ and ⟨*ē*_±*n*_⟩ and the autocorrelation over multiple dominance periods are evident.

This source of serial dependency – comparatively slow fluctuations of *ē* – predicts several qualitative characteristics not described previously and now confirmed by experimental observations. First, sequential correlations are predicted (and observed) to be strictly positive at all lags (next period, one-after-next period, and so on)(**Fig. 6d**). In other words, it predicts that several successive dominance periods are shorter (or longer) than average.

Second, due to the contrast-dependence of autocorrelation time, sequential correlations are predicted (and observed) to increase with image contrast (**Fig. 6d**). The experimentally observed degree of contrast-dependence is broadly consistent with pool sizes between *N* =25 and *N* =40 (black and red curves in **Fig. 2e**). Larger pools with hundreds of bistable variables do not express the observed dependence on contrast (not shown).

Third, for high image contrast, reversal sequences are predicted (and observed) to contain extended episodes with dominance periods that are short, or extended episodes with periods that are long (**Fig. 6c**). When quantified in terms of a ‘burstiness index’, the degree of inhomogeneity in predicted and observed reversal sequences is comparable (supplementary text **Burstiness** and **Fig. 14**).

**Figure 14.**
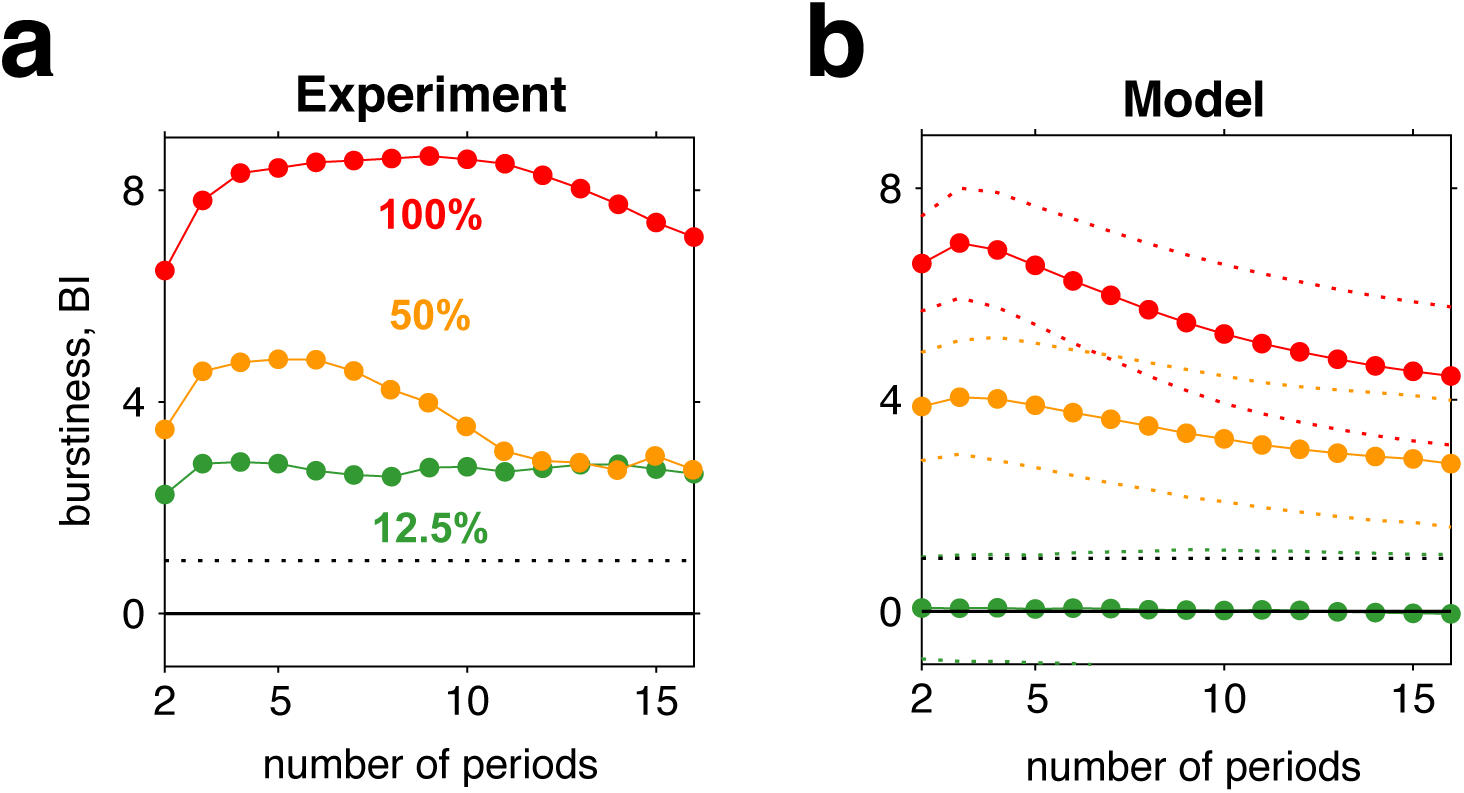
Burstiness of reversal sequences predicted by model and confirmed by experimental observations. **(a)** Burstiness index BI (mean) for *n* successive dominance periods in experimentally observed reversal sequences, for contrasts 12.5% (green), 50% (yellow) and 100% (red). **(b)** Burstiness index for reversal sequences generated by model (mean±std).

Many previous models of binocular rivalry [*e.g*., 59] postulated selective adaptation of competing representations to account for serial dependency. However, selective adaptation is an *opponent* process which ensures *positive* correlations between *different* dominance periods, but *negative* correlations between *same* dominance periods. To demonstrate this point, we fitted such a model to reproduce our experimental observations (*T, c*_*v*_, *γ*_1_, and *cc*_1_) for five image contrasts *c* = *c*′. As expected, the alternative model predicts *negative* correlations *cc*_2_ for *same* dominance periods (**Fig. 6d**, right panel), contrary to what is observed.

## DISCUSSION

We have shown that the universal statistical features of multistable phenomena in perception, as exemplified by binocular rivalry, are reproduced and indeed guaranteed by a particular biophysical mechanism. The explanatory power of this mechanism is extraordinarily high, in that the phenomena explained are considerably more complex (have far more degrees of freedom) than the mechanism itself. It has long been suspected that multistable phenomena in perception may share a similar mechanistic origin. Here we present for first time a possible common mechanism that is biophysically detailed and neurally plausible.

### Continuous inference

The notion that multistable phenomena such as binocular rivalry reflect active exploration of explanatory hypotheses for sensory evidence has a venerable history [5, 44, 65, 118]. The mechanism proposed here is entirely in keeping with that notion: higher-level ‘explanations’ compete for control (‘dominance’) of phenomenal appearance in terms of their correspondence to lower-level ‘evidence’. An ‘explanation’ takes control if its correspondence is sufficiently superior to that of rival ‘explanations’. The greater the superiority, the longer control is retained. Eventually, alternative ‘explanations’ seize control, if only briefly. This manner of operation is also consistent with computational theories of ‘analysis by synthesis’ or ‘hierarchical inference’, although there are many differences in detail [86, 91, 97].

Interacting with an uncertain and volatile world necessitates continuous and concurrent evaluation of sensory evidence and selection of motor action [22, 43]. Multistable phenomena exemplify continuous decision-making without external prompting [17]. Sensory decision-making has been studied extensively, mostly in episodic choice-task, and the neural circuits and activity dynamics underlying episodic decision-making – including representations of potential choices, sensory evidence, and behavioural goals – have been traced in detail [22, 42, 57, 122]. Interestingly, there seems to be substantial overlap between choice representations in decision-making and in multistable situations [17].

### Dynamical mechanism

Two principal *alternatives* have been considered for the dynamical mechanism of perceptual decision-making: drift-diffusion models [69, 98] and recurrent network models [121, 122]. The mechanism proposed here *combines* both alternatives: at its evidence level, sensory information is integrated, over both space and time, by ‘local attractors’ in a discrete version of a drift-diffusion process. At its decision level, the population dynamics of a recurrent network implements a winner-take-all competition between ‘non-local attractors’. Together, the two levels form a ‘nested attractor’ system [17] operating perpetually out of equilibrium.

A recurrent network with strong competition typically ‘normalizes’ individual responses relative to the total response [77]. Divisive normalization is considered a canonical cortical computation [20], for which multiple rationales can be found. Here divisive normalization is augmented by indiscriminate feedforward inhibition. This combination ensures that decision activity rapidly and reliably categorizes *differential* input strength, largely independently of *total* input strength.

Another key feature of the proposed mechanism is that a ‘dominant’ decision pool applies feedback suppression to the associated evidence pool. Selective suppression of evidence for a winning hypothesis features in computational theories of ‘hierarchical inference’ [61, 86, 91, 97], as well as in accounts of multistable perception inspired by such theories [27, 46, 124]. A normative reason for feedback suppression arises during continuous inference in uncertain and volatile environments, where the accumulation of sensory information is ongoing and cannot be restricted to appropriate intervals [117]. Here optimal change detection requires an exponentially rising bias *against* evidence for the most likely state, ensuring that even weak changes are detected, albeit with some delay.

The pivotal feature of the proposed mechanism are pools of bistable variables or ‘local attractors’. Encoding sensory inputs in terms of persistent ‘activations’ of local attractors assemblies (rather than in terms of transient neuronal spikes) creates an intrinsically retentive representation: sites that respond are also sites that retain information (for a limited time). Our results are consistent with a few tens of bistable variables in each pool. In the proposed mechanism, *differential* activity of two pools accumulates evidence *against* the dominant appearance until a threshold is reached and a reversal ensues. This discrete non-equilibrium dynamics may well instantiate a variational principle of inference such as ‘maximum caliber’ [35, 94].

### Emergent features

The components of the proposed mechanism interact to guarantee the statistical features that characterize binocular rivalry and other multistable phenomena. Discretely stochastic accumulation of differential evidence *against* the dominant appearance ensures sensitivity of dominance durations to non-dominant input. It also ensures the constant variability (‘scaling property’) and Gamma-like distribution shape of dominance durations. Due to a highly non-trivial interaction with the competitive decision, discretely stochastic fluctuations of evidence level activity express themselves in a serial dependency of dominance durations. These unexpected dependencies are experimentally confirmed and strongly corroborate the proposed mechanism.

### Previous models

How does the proposed mechanism compare to previous ‘dynamical’ models of multistable phenomena? It is of similar complexity in that it assumes two dynamical levels, one slow (accumulation) and one fast (winner-take-all competition). It differs in reversing their ordering: visual input impinges first on the slow level, which then drives the fast level. If also differs in that stochasticity dominates the slow dynamics [as suggested by 115], not the fast dynamics. However, the most fundamental difference is discreteness (pools of bistable variables), which shapes all key dynamical properties.

### Neural substrate

What might be the neural basis of the bistable variables / ‘local attractors’ proposed here? Ongoing activity in sensory cortex appears to be low-dimensional, in the sense that the activity of neurons with similar response properties varies concomitantly [“shared variability”, “noise correlations”, 37, 75, 76, 92, 99]. This shared variability reflects the spatial clustering of intracortical connectivity [25, 62, 81, 84, 101] and unfolds over moderately slow time-scales, with characteristic times in the range of 100 *ms* to 500 *ms*) reported in primate and rodent cortex [26, 37, 75, 76, 92, 99].

Possible dynamical origins of shared and moderately slow variability have been studied extensively in theory and simulation [for reviews, see 47, 58, 77]. Networks with weakly clustered connectivity (*e.g*., 3% rewiring) can express a metastable attractor dynamics with moderately long time-scales [36, 67, 101, 103]. In a metastable dynamics, individual (connectivity-defined) clusters transition spontaneously between distinct and quasi-stationary activity levels (‘attractor states’) [109, 113].

Evidence for metastable attractor dynamics in cortical activity is accumulating steadily [37, 73–76, 99]. Distinct activity states with exponentially distributed durations have been reported in sensory cortex [37, 75], consistent with noise-driven escape transitions [36, 47]. And several reports are consistent with external input modulating cortical activity mostly indirectly, *via* the rate of state transitions [21, 37, 38, 75, 76].

The proposed mechanism assumes bistable variables with noise-driven escape transitions, with transition rates modulated exponentially by external synaptic drive. Following previous work [19, 36, 47], we show this to be an accurate reduction of the population dynamics of metastable networks of spiking neurons.

As multistable phenonema and their characteristics are ubiquitous in visual, auditory and tactile perception, the mechanism we propose may form a general part of sensory processing. It bridges neural, perceptual and normative levels of description and potentially offers a ‘comprehensive task-performing model’ [56] for sensory decision-making.

## METHODS

### Psychophysics

Six practised observers participated in the experiment (4 male, 2 female). Stimuli were displayed on an LCD screen (EIZO ColorEdge CG303W, resolution 2560 × 1600 pixels, viewing distance was 104 cm, single pixel subtended 0.014°, refresh rate 60Hz) and were viewed through a mirror stereoscope, with viewing position being stabilized by chin and head rests. Display luminance was gamma-corrected and average luminance was 50 *cd/m*^2^.

Two grayscale circular orthogonally-oriented gratings (+45° and −45°) were presented foveally to each eye. Gratings had diameter of 1.6°, spatial period 2 *cyc/deg*. To avoid a sharp edge, grating contrast was modulated with Gaussian envelope (starting inner radius 0.6°, *σ* = 0.2°). Tilt and phase of gratings was randomized for each block. Five contrast levels were used: 6.25%, 12.5%, 25%, 50%, and 100%. Contrast of each grating was systematically manipulated, so that each contrast pair was presented in two blocks (50 blocks in total). Blocks were 120 *s* long and separated by a compulsory one-minute break. Observers reported on the tilt of the visible grating by continuously pressing one of two arrow keys. They were instructed to press only during exclusive visibility of one of the gratings, so that mixed percepts were indicated by neither key being pressed (25% of total presentation time). To facilitate binocular fusion, gratings were surrounded by a dichoptically presented square frame (outer size 9.8°, inner size 2.8°).

Dominance periods of ‘clear visibility’ were extracted in sequence from the final 90 *s* of each block and the mean linear trend was subtracted from all values. Values from the initial 30 *s* were discarded. To make comparable the dominance periods of different observers, values were rescaled by the ratio of the all-condition-all-observer average (2.5 *s*) and the all-condition average of each observer (2.5 ± 1.3 *s*). Finally, dominance periods from symmetric conditions (*c*_*left*_, *c*_*right*_) were combined into a single category (*c*_*dom*_, *c*_*sup*_), where *c*_*dom*_ (*c*_*sup*_) was the contrast viewed by the dominant (suppressed) eye.

For the dominance periods *T* observed in each condition, first, second, and third central moments were computed, as well coefficient of variation *c*_*v*_ and skewness *γ*_1_ relative to coefficient of variation:

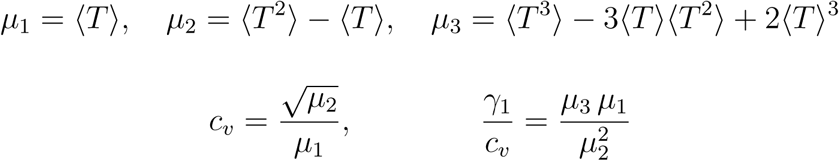

Coefficients of sequential correlations were computed from pairs of periods (*T*_*i*_, *T*_*j*_) with opposite dominance (first and next: ‘lag’ *j* − *i* = 1), pairs of periods with same dominance (first and next but one: ‘lag’ *j* − *i* = 2), and so on,

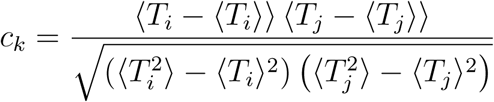

where ⟨*T*⟩ and ⟨*T* ^2^⟩ are mean duration and mean square duration, respectively.

To analyze ‘burstiness’, we adapted a statistical measure used in neurophysiology [24]. First, sequences of dominance periods were divided into all possible subsets of *k* ∈ {2, 3, …, 16} successive periods and mean durations computed for each subset. Second, heterogeneity was assessed by computing, for each size *k*, the coefficient of variation *c*_*v*_ over mean durations, compared to the mean and variance of the corresponding coefficient of variation for randomly shuffled sequences of dominance periods. Specifically, a ‘burstiness index’ was defined for each subset size *k* as

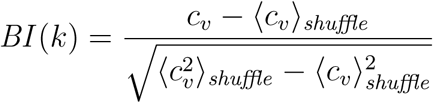

where *c*_*v*_ was the coefficient of variation over subsets of size *k* and where ⟨*c*_*v*_⟩_*shuffle*_ and 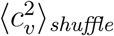 were, respectively, mean and mean square of the coefficients of variation from shuffled sequences.

### Model

The proposed mechanism for binocular rivalry dynamics relies on discretely stochastic processes (‘birth-death’ or generalized Ehrenfest processes). Bistable variables *x* ∈ {0, 1} transition between active and inactive states with time-varying Poisson rates *ν*^+^(*t*) (activation) and *ν*^−^(*t*) (inactivation). Two ‘evidence pools’ of *N* such variables, *E* and *E*′, represent two kinds visual evidence (*e.g*., for two visual orientations), whereas two ‘decision pools’, *R* and *R*′, represent alternative perceptual hypotheses (*e.g*., two grating patterns) (**Fig. 10**). Thus, instantaneous dynamical state is represented by four active counts *n*_*e*_, *n*_*e*′_, *n*_*r*_, *n*_*r*′_ ∈ [0, *N*] or, equivalently, by four active fractions *e, e*′, *r, r*′ ∈ [0, 1].

The development of pool activity over time is described by a master equation for probability *P*_*n*_(*t*) of the number *n*(*t*) ∈ [0, *N*] active variables.

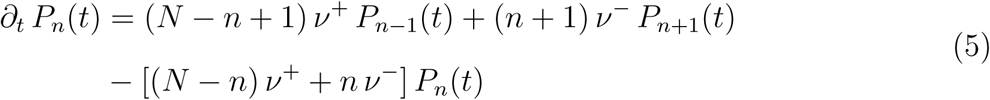

For constant *ν*^±^, the distribution *P*_*n*_(*t*) is binomial at all times [49, 116]. The time-development of the number of active units *n*_*X*_(*t*) in pool *X* is an inhomogeneous Ehrenfest process and corrsponds to the count of activations, minus the count of deactivations,

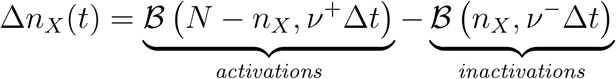

where *ℬ* (*n, ν*Δ*t*) is a discrete random variable drawn from a binomial distribution with trial number *n* and success probability *ν*Δ*t*.

All variables of a pool have identical transition rates, which depend exponentially on the ‘potential difference’ Δ*u* = *u* + *u*^0^ between states, with a input-dependent component *u* and a baseline component *u*_0_:

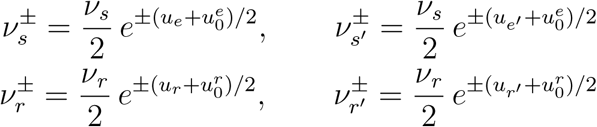

where *ν*_*e*_ and *ν*_*r*_ are baseline rates and 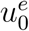 and 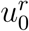 baseline components. The input-dependent components of effective potentials are modulated linearly by synaptic couplings

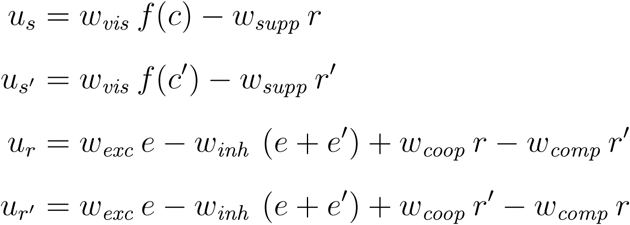

Visual inputs are *I* = *f* (*c*) and *I*′ = *f* (*c*′), respectively, where 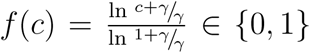 is a monotonically increasing, logarithmic function of image contrast *c* ∈ [0, 1], with parameter *γ*.

### Degrees of freedom

The proposed mechanism has twelve (12) independent parameters: pool size, 6 synaptic couplings, 2 baseline rates, 2 baseline potentials, 1 contrast non-linearity.

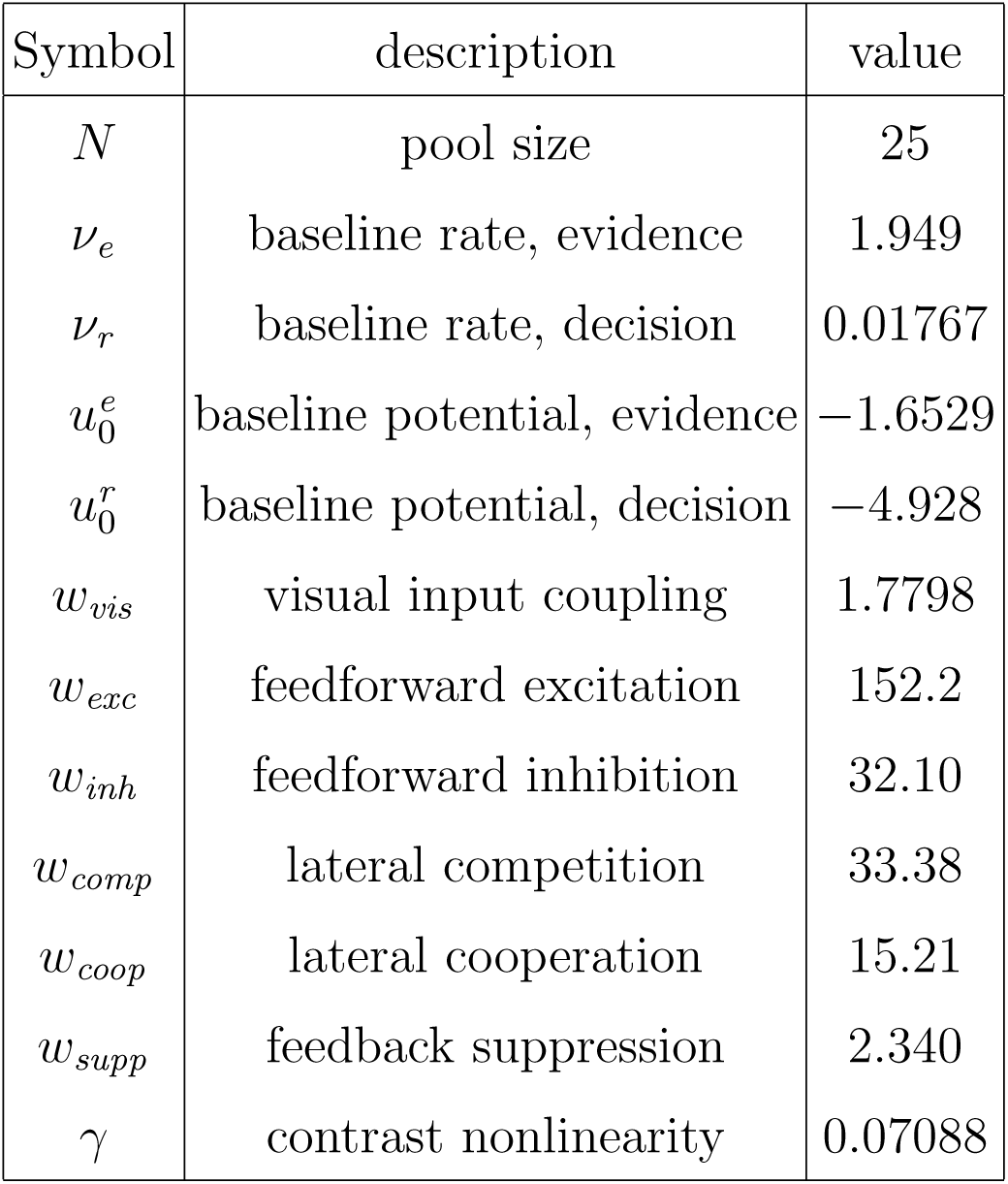

| Symbol | description | value |
| --- | --- | --- |
| $N$ | pool size | 25 |
| $\nu_e$ | baseline rate, evidence | 1.949 |
| $\nu_r$ | baseline rate, decision | 0.01767 |
| $u_0^e$ | baseline potential, evidence | -1.6529 |
| $u_0^r$ | baseline potential, decision | -4.928 |
| $w_{vis}$ | visual input coupling | 1.7798 |
| $w_{exc}$ | feedforward excitation | 152.2 |
| $w_{inh}$ | feedforward inhibition | 32.10 |
| $w_{comp}$ | lateral competition | 33.38 |
| $w_{coop}$ | lateral cooperation | 15.21 |
| $w_{supp}$ | feedback suppression | 2.340 |
| $\gamma$ | contrast nonlinearity | 0.07088 |

### Fitting procedure

The experimental dataset consisted of three 5×5 arrays 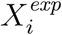 for mean ⟨*T*⟩, coefficient of variation *c*_*v*_, skewness *γ*_1_, plus a scalar correlation coefficient *cc*_1_=0.06, which was the average *cc*_1_ observed over the 5×5 array. In other words, the fitting procedure prescribed contrast-dependencies for distribution moments, but not for correlation coefficients.

The fit error *E*_*fit*_ was computed as a weighted sum of relative errors

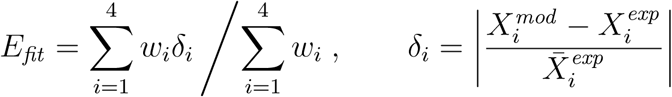

with weighting *w* = [1, 1, 1, 1*/*_4_] emphasizing distribution moments.

Approximately 400 minimization runs were performed, starting from random initial configurations of model parameters. For the optimal parameter set, the resulting fit error for the mean observer dataset was approximately 15%.

To confirm that resulting fit was indeed optimal, and could not be further improved, we studied the behaviour of the fit error in the vicinity of the optimal parameter set. For each parameter *α*_*i*_, 10 values 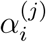 were picked in the direct vicinity of the optimal parameter 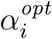. The resulting scatter plot of value pairs 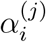 and fit error 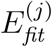 is approximated by a quadratic function. For most parameters the estimated quadratic function is convex, so that the corresponding coefficient of the Hessian matrix associated with the fit error is positive. Also, the estimated extremum of each parabola was close to the corresponding optimal parameter, indicating that the parameter set was indeed optimal.

### Alternative model

As an alternative model [59], a combination of competition, adaptation, and noise was fitted to reproduce ⟨*T*⟩, *c*_*v*_, *γ*_1_, and *c*_1_ for a given image contrast *c* = *c*′. The model comprises four state variables and independent coloured noise,

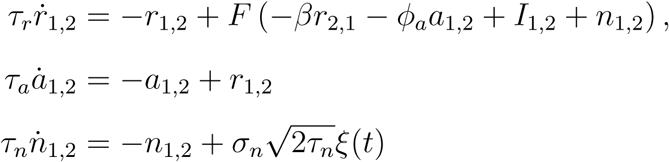

where *F* (*x*) = [1 + exp (− *x/*_*k*_)]^−1^ is a non-linear activation function and *ξ*(*t*) is white noise. Parameters *τ*_*r*_ = 10 *ms, τ*_*n*_ = 100 *ms*, and *k* = 0.1 were fixed. Parameters for competition *β*, adaptation strength *φ*_*a*_, adaptation time-constant *τ*_*a*_, and noise amplitude *σ*_*n*_ were fitted separately for each image contrast.

### Spiking network simulation

To illustrate a possible neural realization of ‘local attractors’, we simulated a competitive network with eight (8) identical assemblies of excitatory and inhibitory neurons, which collectively expresses a spontaneous and metastable dynamics [74]. One assembly (denoted as ‘foreground’) comprised 125 excitatory leaky-integrate-and-fire neurons, which were weakly coupled to the 875 excitatory neurons of the other assemblies (denoted as ‘background’), as well as 250 inhibitory neurons. Note that background assemblies are not strictly necessary and included only for the sake of verisimilitude. The connection probability between any two neurons was *c* = 80%. Excitatory synaptic efficacy between ‘foreground’ neurons, ‘background’ neurons’, and between the two was *J*_fore_ = 0.618*mV, J*_back_ = 0.438*mV*, and *J*_inter_ = 0.402*mV*, respectively. Inhibitory synaptic efficacy was *J*_*I*_ = −1.50*mV* and the efficacy of excitatory synapses onto inhibitory neurons was *J*_*IE*_ = 0.560*mV*. Finally, ‘foreground’ neurons, ‘background neurons’, and ‘inhibitory neurons’ each received independent Poisson spike trains of 2340*Hz*, 2280*Hz*, and 2280*Hz*, respectively. Other settings were as in [74]. As a result of these settings, ‘foreground’ activity transitioned spontaneously between an ‘off’ state of approximately 3 *Hz* and an ‘on’ state of approximately 40 *Hz*.

## ACKNOWLEDGEMENTS

Funding from EU FP7-269459 Coronet, DFG BR 987/3-1 and DFG 987/4-1. The authors are obliged to Andrew Parker for helpful comments.

## SUPPLEMENTARY MATERIAL

### Metastable attractor dynamics

We postulate assemblies or clusters of neurons with recurrent random connectivity as operative units of sensory representations. In our model, such assemblies are reduced to binary variables with Poisson transitions. Our key assumption is that the rates *ν*^±^ of activation and inactivation events are modulated exponentially by synaptic input (Eq. 1):

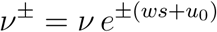

Here we show that these assumptions are a plausible reduction of recurrently connected assemblies of spiking neurons.

Following earlier work, we simulated a competitive network with eight (8) identical assemblies of excitatory and inhibitory neurons (**Fig. 8a**), configured to collectively express a metastable activity dynamics [74]. Here we are interested particularly in the activity dynamics of one excitatory assembly (dubbed ‘foreground’), which expresses two quasi-stable ‘attractor’ states: an ‘on’ state with high activity. In the context of the metastable network, the ‘foreground’ assembly is bistable in that it transitions spontaneously between ‘on’ and ‘off’ states. Such state transitions are noise-driven escape events from an energy well and therefore occur with Poisson-like rates *ν*^+^ (activation) and *ν*^−^ (inactivation). **Fig. 1b** and **Fig. 8b** illustrate this energy landscape for the ‘diffusion limit’ a very large assemblies, where quasi-stable activity levels are *ν*_*fore*_ ≃ 45 *Hz* for the ‘on’ state state and *ν*_*fore*_ ≃ 4 *Hz* for the ‘off’ state. For small assemblies with fewer neurons, the difference between ‘on’ and ‘off’ states is less pronounced.

To establish the dependence of transition rates on external input to the ‘foreground’ assembly, we stepped external input rate Δ*ν*_*ext*_ between two values selected from a range Δ*ν*_*ext*_ ∈ [−120 *Hz*, 50 *Hz*] and monitored the resulting spiking activity in individual neurons, as well as activity *ν*_*fore*_ of the entire population (**Fig. 8c**, upper and middle panels). Comparing population activity to a suitable threshold, we identified ‘on’ and ‘off’ states of the ‘foreground’ assembly (**Fig. 8c**, lower panel), as well as the probability of ‘on’ or ‘off’ states at different points in time following a step in Δ*ν*_*ext*_ (**Fig. 8d**). From the hazard rate (temporal derivative of probability) we then estimated the rates *ν*^±^ of state transitions shown in **Fig. 8d**. Transition rates *ν*^±^ vary anti-symmetrically and exponentially with external input Δ*ν*_*ext*_. In the present example, the dependence is well approximated by *ν*^±^≃ 1*Hz* exp (±0.04*s*Δ*ν*_*ext*_ + 1) (**Fig. 8e**, red and blue lines). This Arrhenius-Van’t-Hoff-like dependence of escape rates is a consequence of the approximately linear dependence of relative potential on external input. Escape kinetics is typical for attractor systems and motivates Eq. 1.

### Quality of representation

#### Accumulation of information

A birth-death process – defined as *N* bistable variables with transition rates *ν*_±_=*νe*^±*ws*^, where *ν* is a baseline rate and *w* a coupling constant – accumulates and retains information about input *s*, performing as a ‘leaky integrator’ with a characteristic time scale [17]. Specifically, the value of *s* may be inferred from fractional activity *x*(*t*) at time *t*, if coupling *w* and baseline rate *ν* are known. The inverse variance of the maximum likelihood estimate is given by the Fisher information

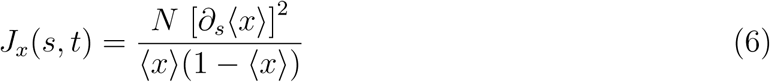

Its value grows with time, approaching *J*_*x*_=*Nw*^2^*/* cosh^2^(*ws/*2) for *t*→∞. For small inputs *s*≃0, the Fisher information increases monotonically as 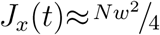 tanh (*νt/*_2_). Surprisingly, the upper bound by 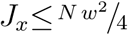 depends linearly on pool size *N*, but quadratically on coupling *w*. Thus, stronger coupling substantially improves encoding accuracy (of input *s*).

The rate at which Fisher information is accumulated by a pool is set by the baseline transition rate *ν*. An initially inactive pool, with *n*_0_=0, accumulates Fisher information at an initial rate of 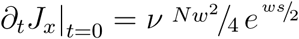. Thus, any desired rate of gaining Fisher information may be obtained by choosing an appropriate value for *ν*. However, unavoidably, after an input *s* has ceased (and was replaced by another), information about *s* is lost at the same rate.

#### Integration of noisy samples

Birth-death processes are able to encode also noisy sensory inputs, capturing much of the information provided. When an initially inactive pool receives an input *s* over time *t*, stochastic activity *n*(*t*) gradually accumulates information about the value of *s*. Normally distributed inputs *s*∈ 𝒩 (*µ, σ*) provide Fisher information *J*_*s*_ = 1/*σ*^2^ about mean *µ*. Pool activity *n*(*t*) accumulates Fisher information *J*_*n*_(*t*) about input mean *µ*, which may be compared to *J*_*s*_. Comparatively small pools with strong coupling (*e.g*., *N* =25, *w*=2.5) readily capture 90% of the information provided (**Fig. 9a**).

Moreover, pools readily permit information from multiple independent inputs to be combined over space and/or time. For example, the combined activity of four pools (*N* =25, *w*=2.5), which receive concurrently four independent samples, captures approximately 80% of the information provided, and a single pool receiving four samples in succession still retains approximately 60% of the information provided (**Fig. 9bc**). In the latter case, retention is compromised by the ‘leaky’ nature of stochastic integration. Whether signals are being integrated over space or time, the retained fraction of information is highest for inputs of moderate and larger variance *σ*^2^ (**Fig. 9bc**). This is because inputs with smaller variance are degraded more severely by the internal noise of a birth-death process (*i.e*., stochastic activations and inactivations).

#### Suitability for inference

Summation of heterogeneous neural responses can be equivalent to Bayesian integration of sensory information [6, 93]. In general, this is the case when response variability is “Poisson-like” and response tuning differs only multiplicatively [70, 71]. We now show that bistable stochastic variables *x*_*i*_(*t*), with heterogeneous transition rates 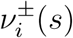, satisfy these conditions, as long as synaptic coupling *w* is uniform.

Assuming initially inactive variables, *x*_*i*_(0) = 0, incremental responses *x*_*i*_(Δ*t*) after a short interval Δ*t* are binomially distributed about mean ⟨*x*_*i*_(Δ*t*)⟩, which is approximately

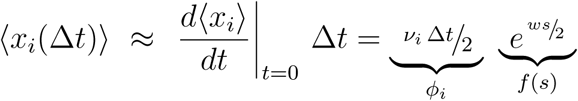

where *ϕ*_*i*_ = *v*_*i*_Δ*t*/2 reflects (possibly heterogeneous) response tuning and *f*(*s*) = *e*^*ws*/2^ represents a common response function which depends only on synaptic coupling *w*. The Fisher information, about *s*, of individual responses is

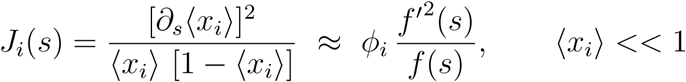

as long as expected activation ⟨*x*_*i*_⟩ is small. The Fisher information of summed responses Σ*i x*_*i*_ is

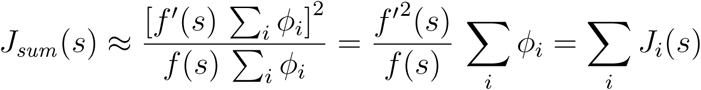

and equals the combined Fisher information of individual responses. Accordingly, the summation of bistable activities with heterogeneous transition rates *ν*_*i*_ optimally integrates information, provided expected activations remain small, ⟨*x*_*i*_⟩<<1, and synaptic coupling *w* is uniform.

### Categorical choice

The ‘biased competition’ circuit proposed here expresses a categorical decision by either raising *r* towards unity (and lowering *r*′ towards zero) or *vice versa*. Here we describe its stochastic steady-state response to constant visual inputs *I*=*f* (*c*) and *I*′=*f* (*c*′) and for arbitrary initial conditions of *e, e*′, *r* and *r*′ (**Fig. 10**). Note that, for purposes of this analysis, evidence activity *e, e*′ was *not* subject to feedback suppression.

The choice is random when the input is ambiguous, *I*≃*I*′, but quickly becomes deterministic with growing input bias |*I*−*I*′|>0. Importantly, the choice is consistently determined by visual input for all initial conditions. The 75% performance level is reached for biases |*I* − *I*′|≈0.04 to 0.06.

Mutual inhibition *w*_*comp*_ controls the width of the ambiguous region around *I*=*I*′, and self-excitation *w*_*coop*_ ensures a categorical decision even for small *I, I*′≃0. The balance between feedforward excitation *w*_*exc*_ and inhibition *w*_*inh*_ eliminates decision failures for all but the largest values of *I, I*′>0.7 and reduces the dependence of sensitivity to differential input |*I*−*I*′| on total input *I*+*I*′.

For particularly high values of input *I, I*′>0.7, no categorical decision is reached and activities of both *r* and *r*′ grow above 0.5. In the full model, such inconclusive outcomes are eliminated by feedback suppression.

### Deterministic dynamics

In the deterministic limit of *N* → ∞, fractional pool activity *x* equals its expectation ⟨*x*⟩ and the relexation dynamics of Eq. 2 becomes

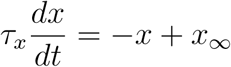

with characteristic time 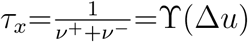 and asymptotic values 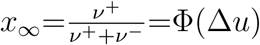, where Δ*u* is the potential difference. Input-dependencies of characteristic time and of asymptotic value follow from Eq. 1:

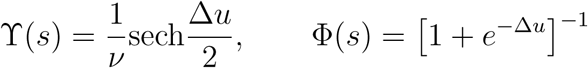

#### Evidence pools

The relaxation dynamics of evidence pools is given by Eq. 2 and Eq. 3’. As shown in the next section, reversals occur when evidence difference |*e* − *e*′| reaches a reversal threshold Δ_*rev*_. For example, a dominance period of evidence *e*′ begins with *e*′_*start*_ =*e*_*start*_ +Δ_*rev*_ and ends when the concurrent habituation of *e*′ and recovery of *e* have inverted the situation to *e*_*end*_ =*e*′_*end*_ +Δ_*rev*_ (**Fig. 11a**). Once the deterministic limit has settled into a limit cycle, all dominance periods start from, and end at, the same evidence levels.

If pool *R*′ has just become dominant, so that *r*′≈1 and *r*≈0, the state-dependent potential differences are

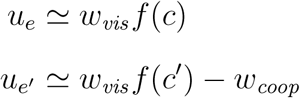

and the deterministic development is

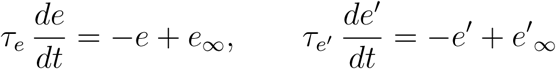

with asymptotic values

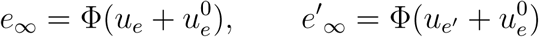

and characteristic times

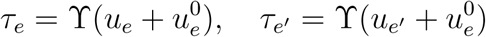

The starting points of the development, *e*_*rev*_ and *e*′_*rev*_ (dashed lines in **Fig. 11ab**), depend mostly on total input *c*+*c*′ and only little on input difference *c*−*c*′. Accordingly, for a given level of total input *c* + *c*′, the situation is governed by the distance between asymptotic evidence levels Δ_∞_=*e*_∞_−*e*′_∞_ and by characteristic times *τ*_*e*_, *τ*_*e*′_.

The dependence on input bias *c*−*c*′ of effective potential Δ*u*_*e*_+*u*_*e*_, characteristic time *τ*_*e*_, and asymptotic value *e*_∞_ is illustrated in **Fig. 11ced**. The potential range of relaxation is *e*_*rev*_ →*e*_∞_ and *e*′_*rev*_ →*e*_∞_, where reversal levels *e*_*rev*_ and *e*′_*rev*_ can be obtained numerically.

Dominance durations depend more sensitively on the slower of the two concurrent processes, as it sets the pace of the combined development. The initial rates *ρ*_*e*_ and *ρ*_*e*′_ after a reversal of the two opponent relaxations

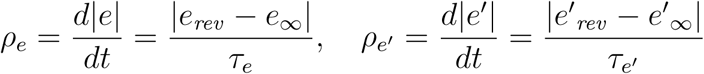

provide a convenient proxy for relative rate. As shown in **Fig. 11f**, when stronger-input evidence *e* dominates, recovery of weaker-input evidence (red up arrow on blue background) is slower than habituation of stronger-input evidence (blue down arrow on blue background). Conversely, when weaker-input evidence *e*′ dominates, recovery of stronger-input evidence (blue up arrow on red background) is slower than habituation of weaker-input evidence (red down arrow on red background). *In short, dominance durations always depend more sensitively on recovery of the currently non-dominant evidence*.

If the two evidence populations *E, E*′ have equal and opposite potential differences, Δ*u*_*e*_=−Δ*u*_*e*′_, then they also have equal and opposite activation and inactivation rates (Eq. 1)

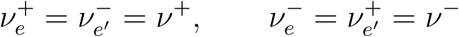

and identical characteristic times *τ*_*e*_ (recovery of *E*) and *τ*_*e*′_ (habituation of *E*′). In this special case, the two processes may be combined and the development of evidence difference Δ*e*=*e*−*e*′ is

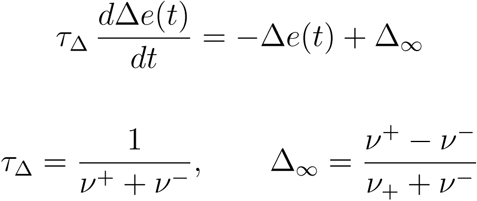

Starting from Δ*e*(0)=−Δ_*rev*_, we consider the first-passage-time of Δ*e*(*t*) through +Δ_*rev*_. If the crossing is certain (*i.e*. when 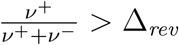), the first-passage-time *T* writes

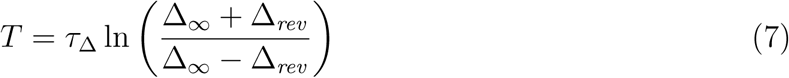

A similar hyperbolic dependence obtains also in all other cases. When the distance between asymptotic levels Δ_∞_ falls below the reversal threshold Δ_*rev*_, dominance durations become infinite and reversals cease.

The hyperbolic dependence of dominance durations, illustrated in **Fig. 11d**, has an interesting implication. Consider the point of equidominance, at which both dominance durations are equal and of moderate duration. Increasing the difference between image contrasts (*e.g*., increasing *c*→*c*+Δ*c* and decreasing *c*′→*c*′−Δ*c* increases Δ_∞_ during the dominance of *e* and decreases it during the dominance of *e*′. Due to the hyperbolic dependence, longer dominance periods lengthen more (*T* →*T* +Δ*T*) than shorter dominance periods shorten (*T* ′→*T* ′−Δ*T* ′), consistent with the contemporary formulation of Levelt III [15].

#### Decision pools

We wish to analyze steady-state conditions for decision pools *R, R*′, as illustrated in **Fig. 5ab**. From Eq. 4, we can write

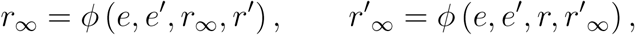

Under certain conditions – in particular, for sufficient self-coupling *w*_*coop*_ – the steady-state equations admit more than one solution: a low-activity fixed point with *r*_∞_≃0, and a high-activity fixed point with *r*_∞_≃1. Importantly, the low-activity fixed point can be destabilized as the activation of the different population changes.

Consider a non-dominant decision pool *R* with fractional activity *r*=*nr/*_*N*_≈0 and its dominant rival pool *R*′ with fractional activity *r*′=*n*_*r*′_/*N*≈1. The steady-state condition then becomes

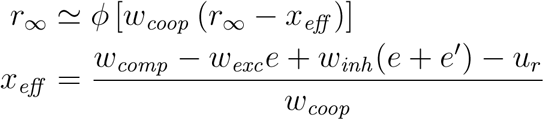

For certain values *x*_*eff*_ ≤*x*_*crit*_, the low-activity fixed point becomes unstable, causing a sudden upward activation of pool *R* and eventually a perceptual reversal. We call the steady-state *r*_∞_ at the point of disappearance *r*_*crit*_.

We can now define a threshold Δ_*rev*_ in terms of the value of evidence bias Δ*e*=*e*−*e*′ which ensures that *x*_*eff*_ ≥*x*_*crit*_ :

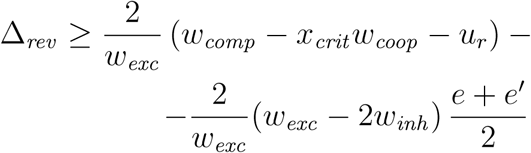

We find that the threshold value Δ_*rev*_ decreases linearly with average evidence 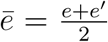, so that higher evidence activity necessarily entails lower thresholds (dashed red line in **Fig. 5c**).

For *w*_*coop*_ = 15.21, we find *x*_*crit*_ = 0.24006, *r*_*crit*_ = 0.0708, and Δ_*rev*_ = 0.4554−1.1564 *ē*.

#### Potential landscape

In **Fig. 5b**, we illustrate the steady-state condition *r*_∞_ = *ϕ* [*w*_*coop*_ (*r*_∞_ − *x*_*eff*_)] in terms of an effective potential landscape *U* (*x*). The functional form of this landscape was be obtained by integrating ‘restoring force’ *F* (*x*) over activity *x*:

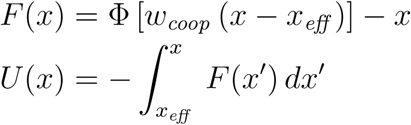

### Stochastic dynamics

#### Poisson-like variability

The discretely stochastic process *x*(*t*) ∈ {0, 1*/*_*N*_, 2*/*_*N*_, …, 1} has a continuously stochastic ‘diffusion limit’, *x*_*diff*_ (*t*), for *N* →∞, with identical mean ⟨*x*_*diff*_⟩= ⟨*x*⟩ and variance ⟨*x*_*diff*_ ^2^⟩− ⟨*x*_*diff*_⟩ ^2^=⟨*x*^2^⟩ −⟨*x*⟩ ^2^. This diffusion limit is a Cox-Ingersoll process and its dynamical equation

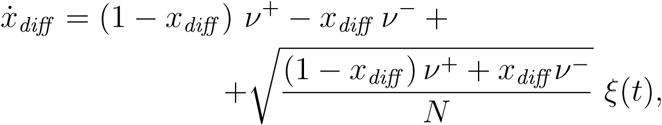

where *ξ*(*t*) is white noise, reveals that its increments 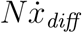 (and thus also the incremenents of the original discrete process) exhibit Poisson-like variability. Specifically, in the low-activity regime, *x*_*diff*_ <<1, both mean and variance of increments approximate activation rate *Nν*^+^:

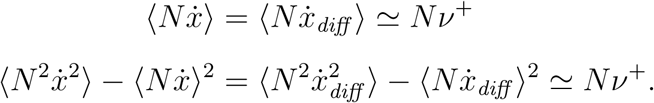

#### Gamma-distributed first-passage times

When the input to a pool of bistable variables undergoes a step-change, the active fraction *x*(*t*) transitions stochastically between old and new steady-states, 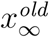 and 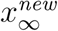 (set by old and new input values, respectively). The time that elapses until fractional activity crosses an intermediate ‘threshold’ level *θ* 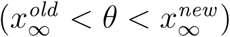 is termed a ‘first-passage time’. In a low-threshold regime, birth-death processes exhibit a particular and highly unusual distribution of first-passage times.

Specifically, the distribution of first-passage times assumes a characteristic, Gamma-like shape for a wide range of value triplets 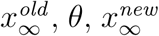 [18]: skewness *γ*_1_ takes a stereotypical value *γ*_1_≃2*c*_*v*_, the coefficient of variation *c*_*v*_ remains constant (as long as the distance between 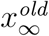 and *θ* remains the same), whereas the distribution mean may assume widely different values. This Gamma-like distribution shape is maintained even when shared input changes during the transition (*e.g*., when bistable variables are coupled to each other) [18].

Importantly, only a birth-death process (*e.g*., a pool of bistable variables) guarantees a Gamma-like distribution of first-passage times under different input conditions [19]. Many other discretely stochastic processes (*e.g*., Poisson process) and continuously stochastic processes (*e.g*., Wiener, Ornstein-Uhlenbeck, Cox-Ingersoll) produce inverse Gaussian distributions with *γ*_1_≃3*c*_*v*_. Models combining competition, adaptation, and noise can produce Gamma-like distributions, but require different parameter values for every input condition (see **Methods:** *Alternative model*).

#### Scaling property

In the present model, first-passage times reflect the concurrent developments of two opponent birth-death processes (pools of *N* =25 binary variables). Dominance periods begin with newly newly non-dominant evidence *e well below* newly dominant evidence *e*′, Δ*e*=*e* − *e*′≃−Δ_*rev*_, and end with the former *well above* the latter, Δ*e*≃+Δ_*rev*_ (**Fig. 12a**). The combination of two small pools with *N* =25 approximates a single large pool with *N* =50. When image contrast changes, distribution shape remains nearly the same, with a coefficient of variation *c*_*v*_≃0.6 and a Gamma-like skewness *γ*_1_≃2, even though mean *µ* of first-passage times changes substantially (**Fig. 12b**).

This ‘scaling property’ (preservation of distribution shape) is owed to the Poisson-like variability of birth-death processes (see above, **Fig. 12c**). Poisson-like variability implies that accumulation rate *µ* and dispersion rate *σ*^2^ are proportional, *µ* ∼ *σ*^2^. This proportionality ensures that activity at threshold disperses equally widely for different accumulation rates (*i.e*., for different input strengths), preserving the shape of first-passage-time distributions [19].

### Characteristic times

As mentioned previously, the characteristic times of pools of bistable variables are not fixed but vary with input (Eq. 2). In our model, the characteristic times of evidence activities lengthen with increasing input contrast and shorten with feedback suppression (**Fig. 13a**). Characteristic times are reflected also in the temporal autocorrelation, which averages over periods of dominance and non-dominance alike. Autocorrelation times lengthen with increasing input contrast, both in absolute terms and relative to the average dominance duration (**Fig. 13b**).

Importantly, the autocorrelation time of *mean* evidence activity *ē*=(*e*+*e*′)*/*2 is even longer, particularly for high input contrast (**Fig. 13c**). The reason is that spontaneous fluctuations of *ē* are constrained not only by birth-death dynamics, but additionally by the reversal dynamics that keeps evidence activities *e* and *e*′ close together (*i.e*., within reversal threshold Δ_*rev*_). As a result, the characteristic timescale of spontaneous fluctuations of *ē* lengthens with input contrast. The amplitude of such fluctuations also grows with contrast (not shown).

The slow fluctuations of *ē* induce mirror-image fluctuations of reversal threshold Δ_*rev*_ and thus are responsible for the serial dependency of reversal sequences (see **Deterministic dynamics:** *Decision pools*).

### Burstiness

The proposed mechanism predicts that reversal sequences include episodes with several successive short (or long) dominance periods. It further predicts that this inhomogeneity increases with image contrast. Such an inhomogeneity may be quantified in terms of a ‘burstiness index’ (BI), which compares the variability of the mean for sets of *n* successive periods to the expected variability for randomly shuffled reversal sequences. In both model and experimental observations, this index rises far above chance (over broad range of *n*) for high image contrast (**Fig. 14**). The degree of inhomogeneity expressed by the model at high image contrast is comparable to that observed experimentally, even though the model was neither designed nor fitted to reproduce non-stationary aspect of reversal dynamics. This correspondence between model and experimental observation compellingly corroborates the proposed mechanism.

